# GSK3 Controls Migration of the Neural Crest Lineage

**DOI:** 10.1101/243170

**Authors:** Sandra G. Gonzalez Malagon, Anna M. Lopez Muñoz, Daniel Doro, Triòna G. Bolger, Evon Poon, Elizabeth R. Tucker, Hadeel Adel Al-Lami, Matthias Krause, Christopher Phiel, Louis Chesler, Karen J. Liu

## Abstract

Migration of the neural crest lineage is critical to its physiological function. Mechanisms controlling neural crest migration are comparatively unknown, due to difficulties accessing this cell population *in vivo*. Here, we uncover novel requirements of glycogen synthase kinase 3 (GSK3) in regulating the neural crest. We demonstrate that GSK3 is tyrosine phosphorylated (pY) in neural crest cells and that this activation depends on anaplastic lymphoma kinase (ALK), a protein associated with neuroblastoma. Consistent with this, neuroblastoma cells with pathologically increased ALK activity express high levels of pY-GSK3 and migration of these cells can be inhibited by GSK3 or ALK blockade. In normal neural crest cells, loss of GSK3 leads to increased pFAK and misregulation of Rac1 and lamellipodin, key regulators of cell migration. Genetic reduction of GSK-3 results in failure of migration. All together, this work identifies a role for GSK3 in cell migration during neural crest development and cancer.

---

The neural crest is a vertebrate-specific, motile population of cells born at the junction of the neural and non-neural ectoderm. This lineage has contributed to our understanding of cellular behaviours, such as contact inhibition of locomotion^1^. It is the origin of many cell types found throughout the organism, including melanocytes, peripheral neurons, cardiac outflow tract, and the craniofacial skeleton. Recent reports have highlighted the importance of neural crest cells: their stem-like capacity, their ability to reprogram, to become cancerous, and to drive vertebrate evolution^2–5^. The highly migratory activity of these cells is critical to their *in vivo* function, not only are their ultimate tissue descendants widespread in the organism, but failure to regulate migration and differentiation in the correct locations is associated with diseases like neuroblastoma^6–8^. Despite its importance, the specific mechanisms underlying this migratory activity and its control are poorly understood.

In our previous work, we demonstrated a critical role for the pleiotropic kinase GSK3 in craniofacial development^9^; therefore, we sought to understand the regulation of GSK3 in neural crest cells, which are integral to most of the craniofacial structures. In vertebrates, the serine/threonine kinase GSK3 is encoded by two paralogous genes, *GSK3α* and *GSK3β*, which are nearly identical throughout their kinase domains^10,11^, and have over 100 predicted substrates^11,12^. The effect of GSK3 phosphorylation is substrate dependent and variable, ranging from control of protein degradation (β-catenin, MYC) and localization (NFAT), to trafficking (amyloid precursor protein) and cleavage (Gli)^11^.

Given the seemingly ubiquitous expression of GSK3 it is not surprising that fine-tuning of this kinase activity is very complex and not well understood, especially *in vivo*. GSK3 can be inhibited by N-terminal serine phosphorylations on both GSK3α (serine-21/S21) and GSK3β (serine-9/S9). These serines are targeted by kinases such as protein kinase A (PKA) and protein kinase B (PKB/AKT) and phosphorylation at these serines prevents GSK3 binding to its substrates^13^. However, interestingly, mouse mutants carrying non-phosphorylatable alanine substitutions at these sites (*GSK3α^S21A/S21A^; GSK3β^S9A/S9A^*) have no obvious developmental phenotypes and are fertile^13^. Using this rigorous approach, functional defects in these animals were limited to regulation of glycogen synthase activity in response to insulin^13^. This evidence that regulation of GSK3 via PKB-dependent serine phosphorylation is dispensible for embryogenesis raises the possibility that additional regulatory mechanisms may be important for GSK3 activity *in utero*. Indeed, inhibition of GSK3 activity by Wnt ligands is serine independent. Instead, inhibition appears to occur via sequestration of GSK3 in response to ligand^14^. This suggests that there exist distinct pools of GSK3 within the cell which are poised for activity.

One positive regulatory mechanism may be via phosphorylation of conserved tyrosine residues (GSK3α-Y279, GSK3β-Y216) which can change the target selectivity of GSK3^15^. While these can be autophosphorylations^16,17^, a recent computational study identified GSK3a as a specific substrate for the kinase ALK, which is often pathologically increased in neuroblastoma cells^18^. However, these *in silico* predictions had not been validated *in vivo* and there had been no studies describing the tissue localisation of this phosphorylation. Given the importance of ALK in pathogenesis of neural crest-derived cancers, we considered the possibility that phosphorylation of these tyrosines might be an important mechanism for ALK-dependent tuning of GSK3 activity, which should be specific to the neural crest lineage. Indeed, we identified specific expression of ALK and phospho-tyrosine GSK3 in the delaminating neural crest cells, as well as a requirement for GSK3 in the control of neural crest and neuroblastoma cell migration.

As noted above, GSK3 is notoriously promiscuous, with many predicted substrates. This has led to reported targets involved in a broad range of biological processes such as senescence, cell proliferation, axonal outgrowth and signaling. GSK3 is also considered a prime therapeutic target in diverse diseases such as diabetes, depression, neurodegeneration, cancer and retinitis pigmentosa^19,20^. As a consequence it is thought that regulation of GSK3 target selection is very context dependent.

Even focusing on the neural crest lineage, GSK3 is thought to have multiple sequential roles beginning with a requirement in patterning of the dorsal axis^21–23^. Based primarily on data from chicken and frog, neural crest-specific GSK3 targets include the Wnt effector β-catenin, the metalloprotease ADAM13 and transcription factors snail and twist^24–26^. Wnt dependent inhibition of GSK3 is clearly necessary for β-catenin-mediated transcriptional activation during neural crest induction^27^. GSK3 also has proposed roles in the phosphorylation of Twist and Snail, proteins which can regulate the activity and stability of these targets, thus controlling the onset of the epithelial-mesenchymal transition (EMT)^24^. Concurrently, GSK3 interactions with ADAM13 are proposed to be crucial for delamination and entry into the EMT^25^. Given the variety of substrates and the precise timing of development, there is no doubt that GSK3 activity must to be dynamically regulated during neural crest development. However, as noted, mice lacking the inhibitory phosphorylation sites at S21/S9-GSK3α/β have normal craniofacial development and are viable^13^. Therefore, we focus on positive regulation of GSK3 via activating tyrosine phosphorylations.

Here, our analysis uncovers a surprisingly specific activation of GSK3 in neural crest cells as they depart from the neuroepithelium and become migratory mesenchymal cells. This activation is controlled by anaplastic lymphoma kinase, which has been implicated in neuroblastoma and other cancers. Genetic and pharmacological loss of GSK3 activity leads to cytoskeletal changes in migratory neural crest cells as well as in neuroblastoma, raising the possibility that control of GSK3 is a rapid and reversible mechanism for controlling cell migration dynamics in the neural crest lineage.

## Results

### GSK3 is expressed, and specifically tyrosine phosphorylated, in migrating neural crest cells

In the embryo, neural crest cells are induced at the neuroepithelial border, subsequently delaminating and becoming migratory. To confirm whether GSK3 was preferentially enriched during neural crest cell migration, we examined mRNA and reporter gene expression for both paralogous genes in frog and mouse, and found that these genes were indeed expressed in migratory neural crest (Figure 1A-H). We noted in particular the enrichment of GSK3 expression in the neural plate stages, at the border of the neural plate (NPB) (Figure 1E, E’, G, G’) and later on in the migratory neural crest cell population, including that destined for the first branchial arch (marked by 1, Figure 1B, D, F, H). The close protein similarity between GSK3 in frog and mouse, as well as the similar expression patterns, suggests that GSK3 activity may play a conserved role in the vertebrate neural crest.

**Figure 1.**
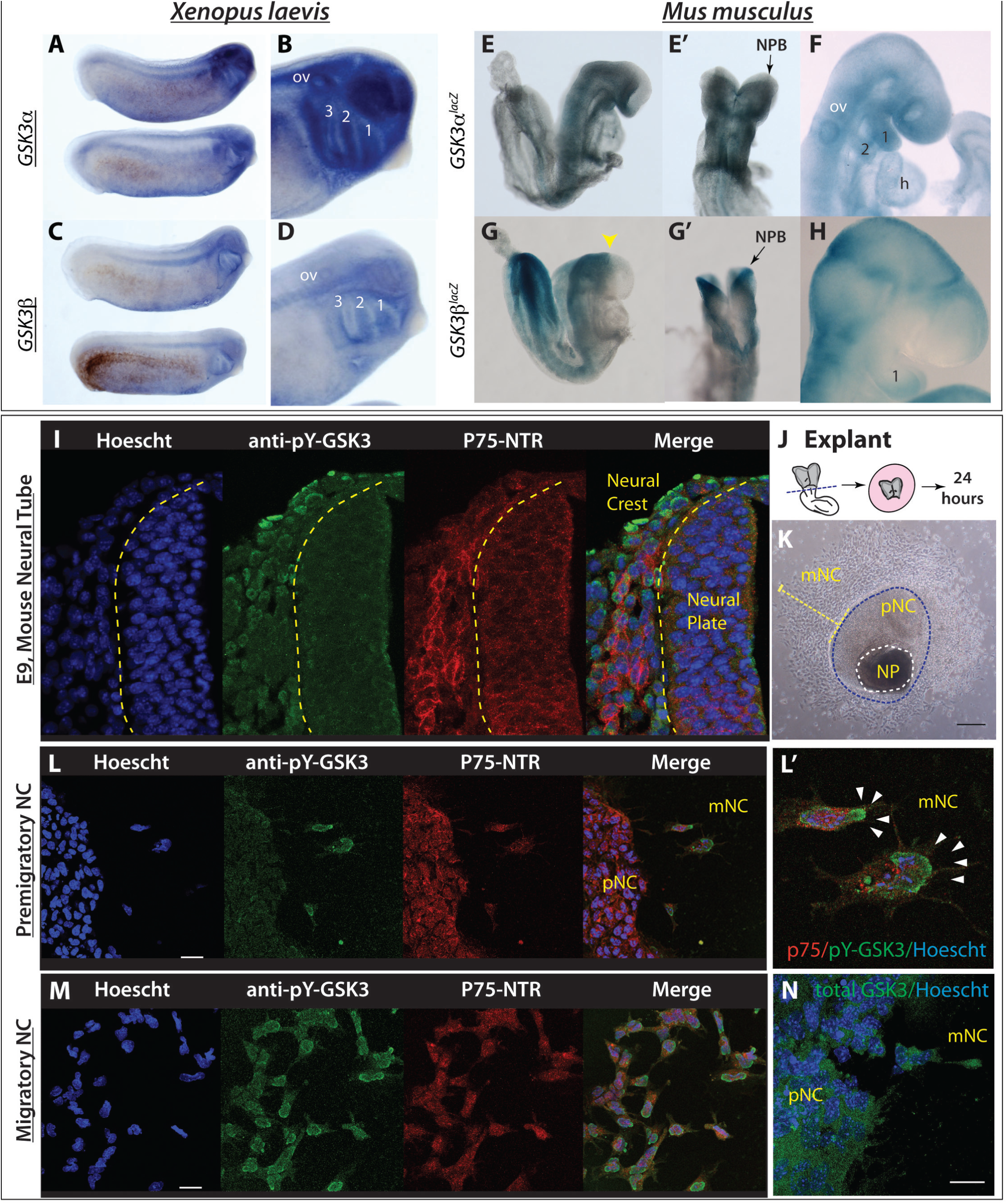
GSK3 genes are expressed during neural crest migration in the frog *Xenopus laevis* and mouse *Mus musculus*. (A-B) mRNA *in situ* hybridization for *Gsk3α* in *X. laevis* at st 25 (A) show expression in the pharyngeal pouches, brain, spinal cord and eye vesicle (B). (C-D) *In situ* hybridization for *Gsk3β* in *X. laevis* at st 25 (C). GSK3β is expressed predominately in the pharyngeal pouches and the spinal cord as well as regions of the brain (D). (E-F). *GSK3α^lacZ^* is expressed in mice during neural crest migration stages. (E-E’) In an e8.5 embryo *GSK3α^lacZ^* is expressed in the cephalic mesenchyme, in the neuroepithelium and in the cephalic neural fold. (F) By E9.5-10, *GSK3α^lacZ^* is highly expressed in the first and second branchial arches (1 and 2) and the frontonasal prominence. (G-H) *GSK3β^lacZ^* is expressed in mice when neural crest is actively migrating. (G-G’) In E8.5 embryos *GSK3β^lacZ^* is mainly expressed in the neuroectoderm, restricted to the prospective hindbrain and some areas in the mesenchyme. (H) At e9.5, *GSK3β^lacZ^* is mainly expressed in BA1 and cranial ganglia, and in the presumptive trigeminal ganglion. **GSK3α/β are phosphorylated at tyrosines Y216/279 during cranial neural crest cell migration**. (I) Transverse cranial section of E9 mouse showing immunoflourescent staining for Hoechst/DNA (blue), pY-GSK3 (green) and p75^NTR^ (neural crest, red). (J) Schematic of E8.5 mouse embryo depicting cranial neural crest (CNC) dissection. (K) Brightfield image of mouse neural crest explant. Neural plates (NP) were dissected from E8.5 embryos and cultured for 24 hours. Two types of cells surround the NP: pre-migratory neural crest cells (pNC) which are epithelial, and migratory neural crest (mNC) scale bar, 250μm. (L) Cells migrating away from the premigratory neural crest begin to express pY-GSK3. Premigratory neural crest (pNC) to the left. All neural crest express p75NTR (red). Note in merge that perinuclear expression of pY-GSK3 is invariably oriented in direction of migration (L’ white arrowheads). (M) Migratory NC cells express pYGSK3 (green) and p75-NTR (red). (L-M) scale bars = 25*μ*M. (N) Expression of total GSK3 is ubiquitous in premigratory and migratory neural crest cells. Scale bar = 25*μ*M.

We were then curious whether GSK3 proteins were activated at specific time points during murine neural crest development. To address this, we used an antibody recognizing a phosphorylated tyrosine in the active site of both GSK3 isoforms (pY279-GSK3α/pY216-GSK3β, referred to hereafter as pY-GSK3). These sites are identical in the two proteins. pY-GSK3 (green) was specifically detected in the cranial neural crest cell population (marked by P75-NTR, red) after emigration from the neural tube (Figure 1I). This was in contrast to more widespread mRNA expression of GSK3α/β seen above (Figure 1A-H). This phosphorylation was also confirmed in a simple *ex vivo* culture system, which allows us to visualize and manipulate specific neural crest populations without complications from surrounding tissues (Figure 1J). In these assays, neural plates from embryonic day 8.5 (E8.5) mouse embryos were explanted and cultured *in vitro*, prior to neural crest migration, allowing subsequent examination of delaminating neural crest cells. By 24 hours of culture, the premigratory neural crest cells (pNC) are spread in an epithelial sheet surrounding the neural plate (NP), with fully migratory cells (mNC) in the outer ring (Figure 1K). Again, we noted that pY-GSK3 is specifically found in neural crest cells just when they delaminate and become mesenchymal (Figure 1L-M). In fully migratory cells, the majority of pY-GSK3 appears perinuclear and is invariably oriented in the direction of migration (Figure 1L-L’). In contrast, staining for total GSK3 appears diffuse and ubiquitous (Figure 1N).

### In vivo loss of GSK3α/β prevents migration of cranial neural crest cells

We then turned to mouse mutants to determine the *in vivo* genetic requirements for *GSK3* in the neural crest. To do this, we generated embryos with a conditional deletion of both *GSK3α* and *GSK3β* genes using a neural crest specific cre-recombinase driver (*Wnt1::cre*^28^). By E9.5, we found widening of the neural plate (Figure 2A, 2E, red bracket), with a medial expansion of *Sox10* expression (Figure 2A, 2E, asterisk). By E10.5, dorsal views revealed an accumulation of *Sox10* positive cells in the brain (Fig 2F, red bracket), suggesting that the cranial neural crest cells have failed to migrate from the neural tube. Complete nulls had a severe disruption of cell migration to the facial prominences, the branchial arches and the cranial ganglia (compare *Sox10* positive/blue in Figure 2C-D to 2G-H). Because we rarely found animals surviving beyond E11.5, we also examined animals carrying a single allele of functional GSK3 (*Wnt1::cre; GSK3α^fl/+^;0^fl/fl^* or *Wnt1::cre; GSK3α^fl/fl^;β^fl/+^*). In both cases, we noted a reduction in *Sox10* positive cells en route to the first branchial arch and an accumulation of positive staining in the neural tube, suggesting that both GSK3 proteins contribute to migration of the neural crest (Supplemental Figure 1, red bracket).

**Figure 2.**
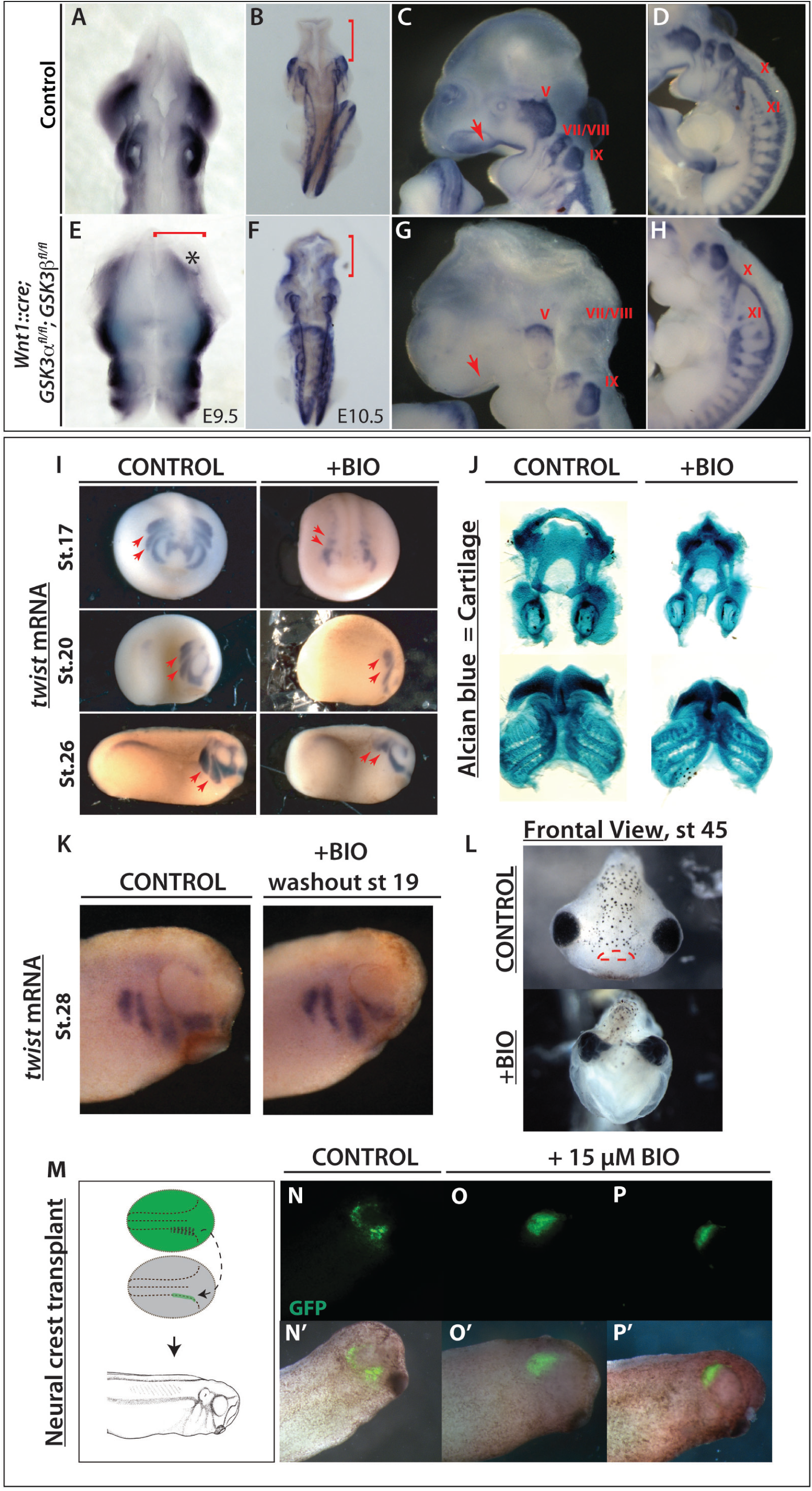
GSK3 is required for neural crest migration *in vivo*. (A to H) mRNA *in situ* hybridization of *Sox10*, which marks migratory neural crest. (A, E) E9.5 mouse embryos. (B-D, F-H) E10.5 mouse embryos. (A-B, E-F) Dorsal views. (C-D, G-H) Lateral views of E10.5 control embryos. (A-B) In control embryos, *Sox10* expression is absent in the brain (B, red bracket) but it is highly expressed along the embryo axis. (E-F) Neural crest specific deletion of GSK3. Dorsal view of E10.5 mutant mouse in which *Sox10* expression has accumulated in the brain (F, red bracket) (C-D) At E10.5, *Sox10* marks cranial neural crest, which has migrated into the facial prominences (C, red arrow) and the cranial ganglia, including the trigeminal ganglia (V) and facial and acoustic nerves (VII/VIII), glossopharyngeal nerve (IX), vagus nerve (X) and the spinal accessory nerve (XI) (D). (G-H) E10.5 mutant mouse lost *Sox10* expression, especially in the facial prominences (G, red arrow) and showed remarkably reduced expression in cranial ganglia and nerves X, XI. The dorsal root ganglia seem to be unaffected. (I) *Twist* expression marks migratory neural crest. BIO treatment from st12.5 results in a loss of *twist* expression at st17 (frontal views, st17). BIO treatment from st12.5 to st19, shows loss of twist expression at st20 and st26 (lateral views). The posterior streams are selectively impaired, red arrows. (J) BIO treatment resulted in a reduction of Alcian blue stained facial cartilages, which are derived from neural crest. (K) *Twist* expression shows that cell migration is regained by stage 28, following washout from the treatment with BIO from st12.5 to st19. (L) Frontal view of a tadpole at stage 45, previously treated with BIO (st.12.5-19). Note narrowing of the head structures and loss of the mouth (marked with red lines in control). (M-P’) GFP labeled neural crest was grafted into a non-labeled embryo at stage 17 and grown to st28. GFP labeled cells in control animals have migrated (N), while those treated with 15*μ*M BIO have not (O-P).

While the timing of the *Wnt1::cre* transgene misses the initiation of neural crest induction, it was still possible that some of the effects seen were due to GSK3 requirements in the pre-migratory neural crest. In order to bypass early effects on the neural crest we turned to pharmacological inhibition of GSK3 using the specific inhibitor BIO (6-bromoindirubin-3’-oxime^29^). Making use of *Xenopus* allowed us to precisely time our manipulations in a well-defined *in vivo* system. Treatments of *Xenopus* embryos confirmed that GSK3 inhibition led to loss of the migration marker *twist1* (Figure 2I). Treatment of *Xenopus* embryos at stage 12.5 confirmed that GSK3 inhibition led to loss of *twist1*. When the embryos were released from treatment at stage 19, we found some recovery of *twist1* expression (Figure 2K). Loss of GSK3 function during the critical stages led to significant changes in face shape as well as a smaller neural crest derived tail fin (Supplemental Figure 2A-C), as well as loss of the neural crest derived facial cartilages (Figure 2J, L). Although the facial cartilages were lost leading to narrowing of the head, the mesodermal cranial muscles are still formed (Supplemental Figure 2D). To confirm that this effect was specifically due to impairment of migration, we transplanted fluorescently labeled neural crest cells into a st.17 embryo and then treated with BIO; these cells did not migrate (Figure 2N-P’). Taken together, this demonstrated a previously under-appreciated role for GSK3 during neural crest cell migration.

### Perturbing GSK3 function prevents migration of cranial neural crest cells

To bypass the neural crest induction stage as well as the embryonic lethality, we turned back to the neural crest explant cultures. We found that dispersion of *Xenopus* cells was inhibited by BIO treatment (Figure 3A-C). Note that in the *Xenopus* explants, the premigratory neural crest population can be specifically dissected independently from the neural plate. We then turned back to mouse neural crest cultures in order to better compare our pharmacological inhibitors to genetic mutants. Treatment with two different specific inhibitors, BIO and CHIR99021 (CHIR^30^) prevented neural crest cell migration similar to that observed in *Xenopus* (Fig 3D-I, Supplemental Movies 1-2). We found that the area covered by the pre-migratory neural crest cells was expanded in treated samples (Supplemental Figure 3A, 3B) with a concurrent decrease in the migratory population (Supplemental Figure 3B), suggesting a defect or delay in the cells emigrating from the leading edge of the neuroepithelium. This was confirmed in genetic mutants, where a complete loss of GSK3 (*Wnt1::cre; GSK3α^fl/fl^;β^fl/fl^*) also led to a decrease in migratory neural crest cells (Figure 3J-L).

**Figure 3.**
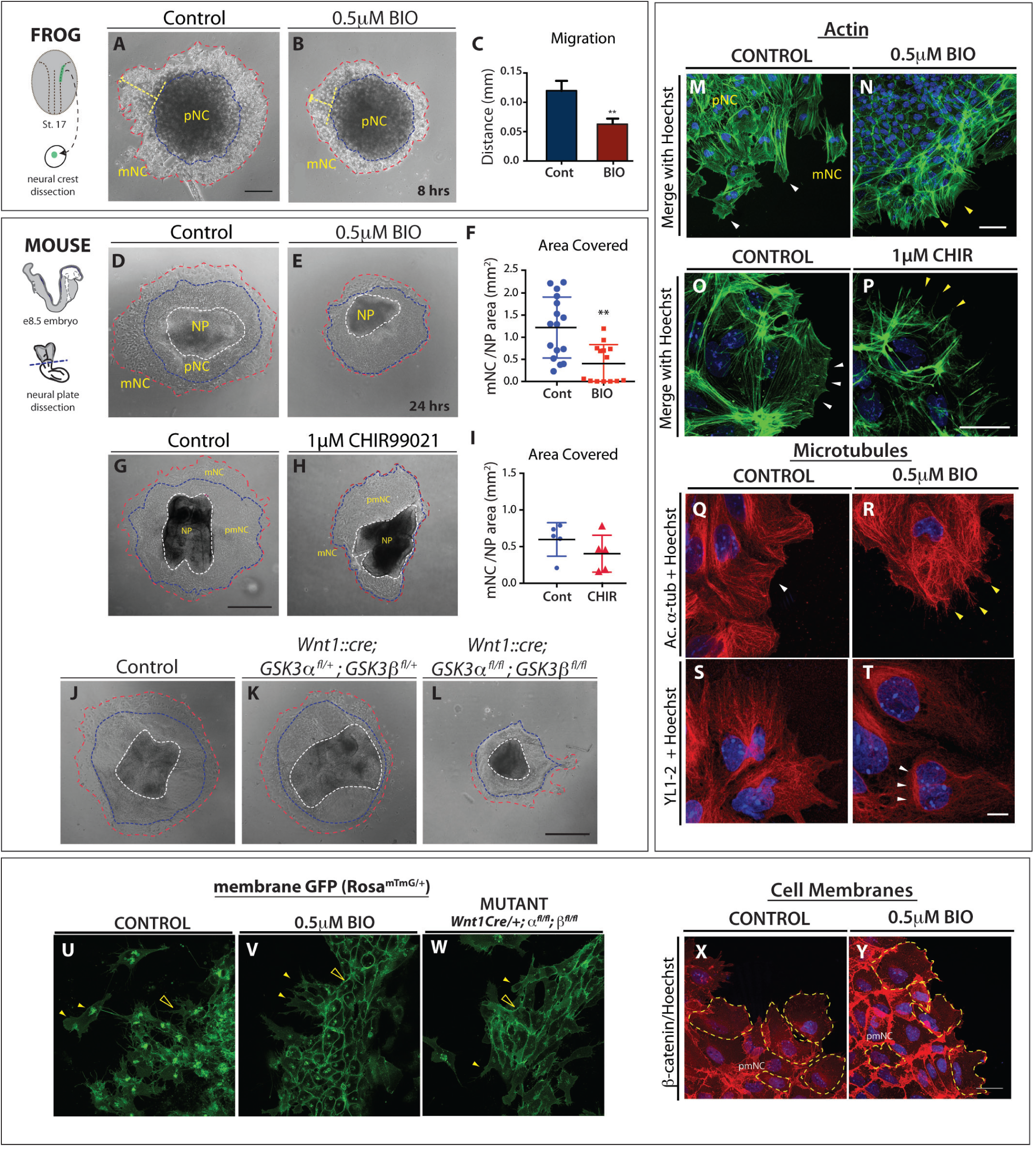
GSK3 activity is required for neural crest cell migration. (A-B) Neural crest cells are dissected from st 17 *Xenopus* embryos. (A) Control explants after 8h in culture. (B) When GSK3 is inhibited (0.5μM BIO) neural crest cells spread significantly less than the controls. (C) Quantification of the distance migrated \*\**p* ≤ *0.05*. (D, G, J) Control mouse explants. (E, H) Explants treated with 0.5*μ*m BIO or 1*μ*M CHIR99021 respectively. Note decrease in area covered by migratory neural crest (F, refers to D, E; I refers to G, H) (J-L) Mouse explants from control (J), *Wnt1::cre; GSK3afl/+; GSK3bfl/+* (K), and *Wnt1::cre; GSK3afl/fl; GSK3bfl/fl* complete mutants (L). Note decrease in area covered in L (red dotted line). (M-P) Phalloidin staining (green) labels filamentous actin in neural crest explants. (M, O) Explants treated with DMSO (control) show accumulation of F-actin in lamellipodia in the leading edge (white arrowheads). Explants treated with (N) 0.5*μ*m BIO and (P) 1μM CHIR lack lamellipodia and only show stress fibers at the cell edge (yellow arrowheads) (Q-T) Microtubules are labeled with acetylated α-tubulin or YL1-2. Note smooth lamellipodial edge in control (Q) and spiky protrusions in BIO treated cells (R). YL1-2 tubulin in control cells is distributed all throughout the cell structure (S); however in BIO treated cells (T), it is mainly found in a perinuclear zone, at the rear of the nucleus. (U) Cranial neural crest explants from control E8.5 embryos carrying membrane GFP in the neural crest lineage. Migrating cells show an elongated morphology and have a dynamic cell-cell contact (see Supplemental Movies 1, 3, 6). Membrane GFP is unstable and intracellular, likely due to recycling of cell membranes. (V) In the BIO treatment, cells remain in contact with adjacent cells and multiple protrusions (yellow arrows). (W) Mutant cells carrying a neural crest specific deletion of GSK3 (*Wnt1Cre/+; GSK3afl/fl; GSK3bfllfl*) lose motility and maintain stable cell-cell contacts and membrane GFP. (X-Y) β-catenin staining in neural crest explants. Note how β-catenin is remarkably stable in BIO treated cells (Y), suggesting increased cell-cell adhesion. (A-L) Scale bar=100*μ*m. (M-Y) Scale bar=25*μ*M.

One possibility was that inhibitory serine phosphorylation of GSK3 is necessary at the leading edge of polarized cells as has been demonstrated in astrocytes and neurons^31–34^. However, mouse mutants carrying non-phosphorylatable S21A/S9A substitutions (*GSK3α^S21A/S21A^; GSK3β^S9A/S9A^*) are viable, suggesting normal neural crest migration^13^. Nevertheless, to confirm this, we observed that neural crest explants from these mice appear normal and retain sensitivity to BIO inhibition, demonstrating that serine phosphorylation of GSK3 is dispensable during neural crest migration (Supplemental Figure 4A-B). Therefore, because of the polarized expression of pY-GSK3 at the leading edge, we decided to focus on the role of GSK3 as the neural crest cells acquire their mesenchymal nature.

### GSK3 inhibition perturbs the cytoskeleton

GSK3 activity has been predicted to control cytoskeletal dynamics in a number of systems. The loss of cell migration in mutant or BIO treated cultures suggested a disruption of cytoskeletal dynamics; therefore, we examined the actin and microtubule arrangements in neural crest cells after inhibition of GSK3 (Figure 3M-T). GSK3 inhibition markedly increased stress fibres and concurrently reduced leading-edge actin (Figure 3M-P), Supplemental Figure5A). Cell shapes at the leading edge were markedly different, with both BIO and CHIR treated cells losing filamentous actin localisation, which is normally at the edge of the cell (see Figure 3M, O, white arrowheads) and generating spiky protrusions (Fig 3N, 3P, yellow arrowheads). Microtubule organization was also disrupted in BIO-treated cells. In normal cells, stabilized microtubules (marked by acetylated tubulin) extend from the centrosome toward the leading edge of the cell (see Figure 3Q). In BIO-treated cells stabilized microtubules appeared to accumulate perinuclearly (Figure 3R, Supplemental Figure 5C). We also examined a marker for unstable microtubules (YL1-2^35^) and found a significant decrease in this population, which also accumulated primarily at the rear of the cell, behind the nucleus (Figure 3S-T, arrowheads and Supplemental Figure 5D). Finally, we examined membrane dynamics in both mutants and BIO-treated explants from mice carrying a genetically labelled membrane GFP (Figure 3U-W). In the control explants, a large proportion of the GFP was internalised within the cell, suggesting recycling of the membrane (Figure 3U, closed arrowhead). Instead, both treated and mutant cells had very strong expression of GFP at the cell membrane (Figure 3V, 3W, open arrowheads). Consistent with these findings, we also found an accumulation of β-catenin at the membrane in BIO-treated explants (Figure 3X-Y), suggesting that loss of GSK3 activity led to extremely stable membrane dynamics compared to controls.

One candidate GSK-3 substrate is focal adhesion kinase (FAK), a well-known regulator of cell motility, which controls focal adhesion dynamics and turnover required for formation of branched actin (as opposed to linear actin). In motile cells, FAK is regulated by an activating tyrosine phosphorylation and a series of inhibitory serine phosphorylations, which are mutually exclusive; of these, S722 is a known GSK3 target^36,37^. Consistent with GSK3 roles in inhibiting FAK, we found that pY-GSK3 (in green) and pY-FAK (in red) were mutually exclusive in the cytoplasm (Figure 4A-B). In addition, we found that migratory neural crest cells ordinarily express active pY-FAK in a halo of puncta that are oriented toward the direction of cell movement, which is toward the right side of all figures (Figure 4C, E, G’). This punctate expression is reminiscent of microspikes, or transient actin-filled filopods which initiate actin nucleation filaments. Percentage of cells containing pFAK at the edge was consistently reduced after GSK3 inhibition (Fig 4J-K) along with a loss of branched actin (Fig 4I) suggesting a loss of lamellipodia-like characteristics. Instead, loss of GSK3 led to a striking accumulation of active pY397-FAK in long extensions indicating persistent focal adhesions (Figure 4F, L-M).

**FIGURE 4.**
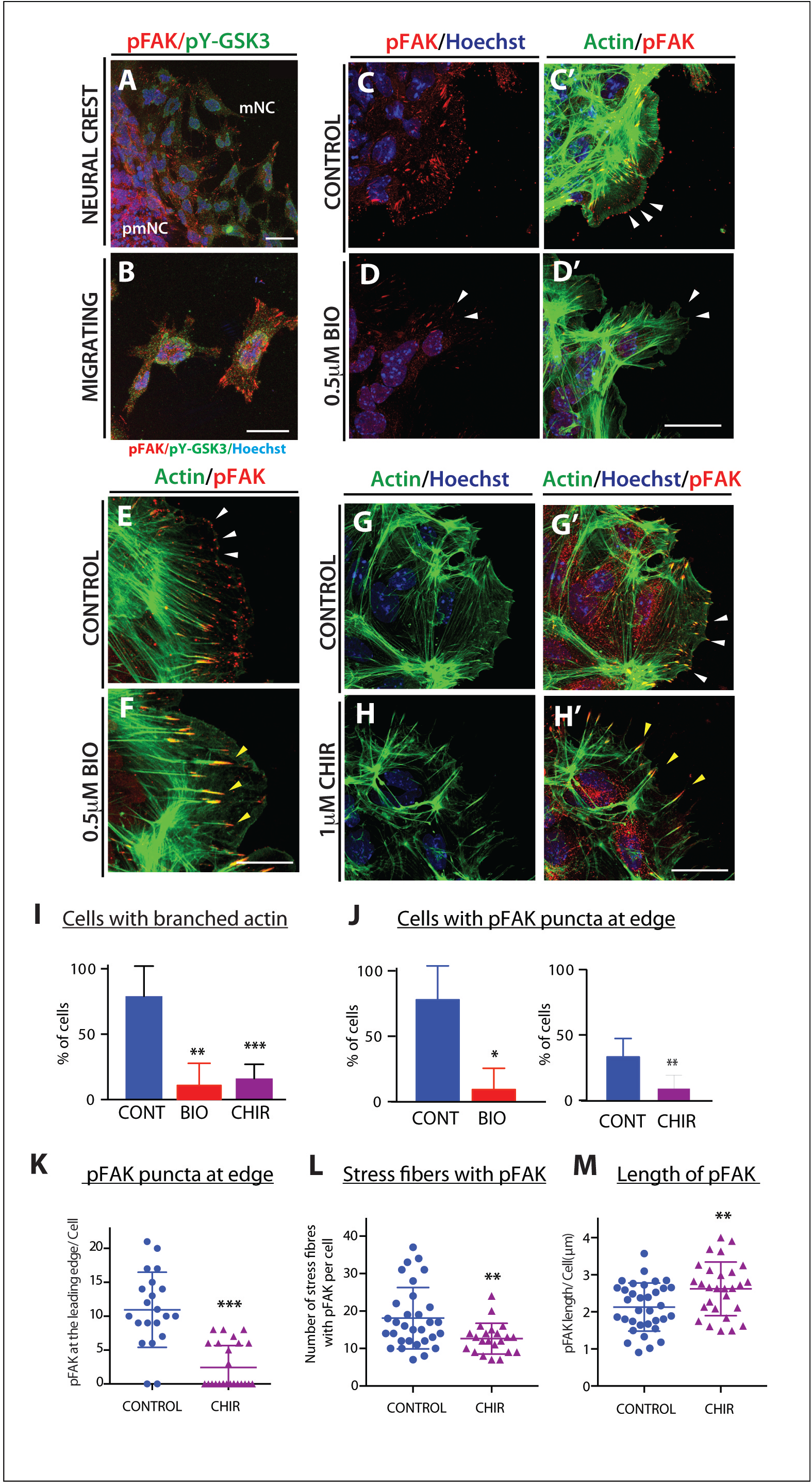
GSK3 allows FAK localisation to establish lamellipodial protrusions at the leading edge in migrating neural crest cells. (A-B) Immunofluorescence for pFAK-Y397 (pFAK, red) and pY-GSK3 (green). (A) In control explants pFAK is found at the leading edge of the delaminating cells and in migrating cells. (B) In migratory neural crest cells, pYGSK3 and pFAK are mutually exclusive. (C-C’) pFAK is found in puncta at the leading edge of the cell co-localizing with lamellipodia. (D-D’) Upon GSK3 inhibition the cells lose pFAK at the leading edge. (E-F) pFAK accumulates at the tips of actin-rich fibres. Note increase in length of pFAK associated with actin upon treatment with BIO (F). (G-H’) Similarly, treatment with 1*μ*M CHIR elicits the same response (white and yellow arrowheads). (A-H) Scale bar=25*μ*M. (I-J) Percentage bar charts representing a significant decrease in cells with branched actin (I) or showing pFAK puncta at the leading edge (J) when GSK3 is inhibited with either BIO or CHIR *\*\*p≤ 0.001 and ***p≤ 0.0001*. (K-M) Dot plots representing the number of pFAK puncta at the edge (K), the number of stress fibers containing pFAK (L) and the length of of pFAK (M) in control and CHIR treated cells; each dot represents one cell. *\*\*p≤ 0.001 and ***p≤ 0.0001*.

FAK is thought to control cytoskeletal dynamics by repressing the function of the small GTPase Rac1^38^. Therefore, the inhibition of FAK activity appears necessary to allow Rac1 activation. With the accumulation of active FAK in our treated cells, we found that Rac1 was now excluded from the leading edge of the migratory neural crest cells (compare Figure 5B to A, arrowheads). Interestingly, we see an increase in Rac1 in nuclei (Figure 5B-C), which could indicate a more direct role for GSK3 in subcellular localisation of Rac1, possibly via phosphorylation of the Rac1 regulator nucleophosmin/B23^39,40^. We saw a concurrent loss of cdc42 localization to the leading edge of the cell (Figure 5D-E). Finally, since FAK and Rac1 can control lamellipodial movement, we examined the localisation of lamellipodin, which regulates neural crest migration via the actin effectors Ena/VASP and Scar/WAVE^41,42^. When GSK3 is inhibited, leading edge localisation of lamellipodin is lost, and surprisingly, lamellipodin also relocalizes to the nucleus (Figure 5F, 5G). As a consequence, treated neural crest cells fail to generate stable fan-shaped lamellipodia (Figure 5H-K, green arrowheads), with approximately 50% of delaminated cells having unstable lamellipodia (Figure 5K). We then turned back to genetic mutants to confirm these phenotypes (Fig 5L-M). In this case, to bypass any complications of GSK3 involvement in neural crest induction, we turned to a tamoxifen inducible Cre (*pCAAG::CreER™*)^43^, allowing temporal deletions upon drug addition. As predicted, tamoxifen induced knockout of *GSK3* led to a loss of the wavelike lamellipodial protrusions, leaving only spiky filopodial movements in neural crest cells (stills shown in Figure 5L-M and Supplemental Movies 3-5). All together, this demonstrates that in the absence of GSK3 activity, neural crest cells make filopodial protrusions at the expense of lamellipodia.

**Figure 5.**
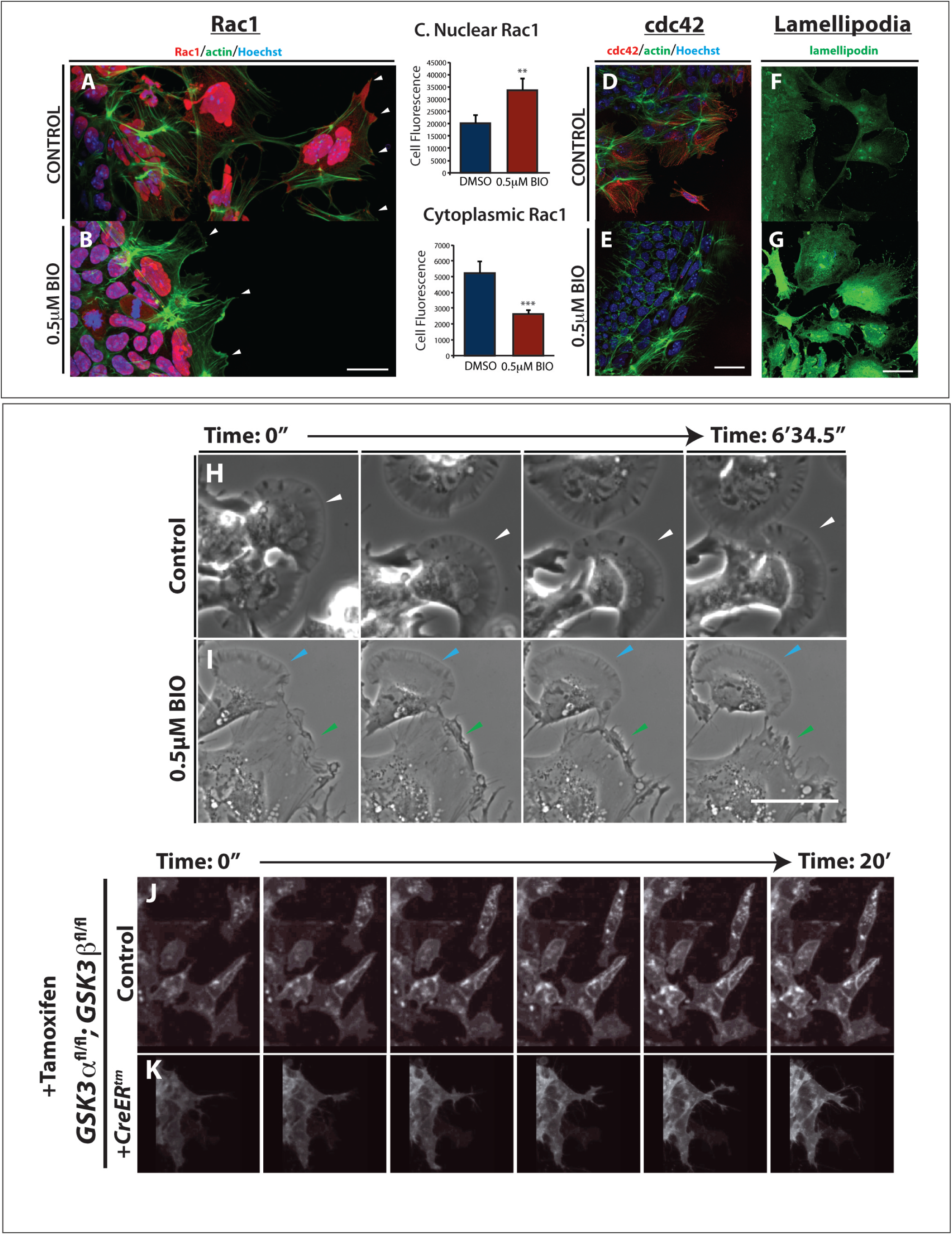
GSK3 is required to establish polarity and to form lamellipodia in migrating neural crest cells. (A-B) GSK3 inhibition increases nuclear RAC1 and reduces cytoplasmic RAC1 in neural crest cells. (A) In control explants RAC1 (red) is high in the nucleus while in the cytoplasm it is enriched at cell protrusions (arrowheads). Actin was labeled with phalloidin (green). (B) In BIO treated explants nuclear RAC1 is high but cytoplasmic staining is lost (white arrowheads). (C) Relative levels of Rac1 fluorescence in the nucleus or the cytoplasm, *\*\*p≤0.001 and ***p≤0.0001* (D-E) GSK3 inhibition reduces expression of cdc42 in neural crest cells. (D) Cdc42 is cytoplasmic and perinuclear in neural crest cells. (E) BIO treated explants lose cdc42 staining. (F-G) In controls, anti-lamellipodin (green) stains the ruffled edge of migrating cells. (G) In contrast, BIO-treated cells show increased total lamellipodin throughout the cell, losing specific localisation at the cell edge. Scale bar, 20μm. (H-I) Brightfield still images from controls (H) or BIO treated samples (I). Control images show cells form fan-shaped stable lamellipodia and ready to migrate away from the cluster (H, white arrowheads). In treated cells, despite some cells form stable lamellipodia as found in controls (I, light blue arrowheads), cells predominantly formed an irregular protrusion that tend to retract (I, green arrowheads) Scale bar, 500μm (See corresponding Supplemental Movies 6, 7) (J) Graph showing the number of cells delaminating from the pre-migratory neural crest cell clusters in two hours. (K) Percentage of delaminating cells which show stable (persistent) lamellipodia or unstable (short-lived) lamellipodia. (L-M) Still images of time-lapse showing control mouse neural crest cells (L, supplemental movies 3, 5) and *pCAAG::creER^tm^; GSK3α^fl/fl^; GSK3β^fl/fl^*; *Rosa^mtmg/+^* deleted cells (M, supplemental movie 4,5). Upon tamoxifen induced knockout of GSK3, the neural crest cells are unable to migrate and the cell edge does not form lamellipodia.

### Anaplastic lymphoma kinase (ALK) is expressed in migratory neural crest cells

Aberrant neural crest development is thought to underlie neuroblastoma, an aggressive paediatric cancer. Activating mutations in ALK contribute to a subset of neuroblastoma cases, correlating with poor prognosis^44–49^. Because we saw specific expression of pY-GSK3 at the leading edge and in migratory neural crest cells, we wondered whether ALK might be responsible for activating GSK3. First, we set out to check whether ALK was expressed during the appropriate stages of neural crest development. Expression of ALK has previously been reported at E10.5, including in the diencephalon and facial ganglia^50^; however, to our knowledge, there has been no analysis performed during key neural crest migration stages. To test this, we performed mRNA *in situ* hybridization from E8.0 onwards (Figure 6A, D). We found that by E8.5, ALK appeared specific to the neural plate border corresponding to active cranial crest migration (Fig 6A). This expression was enriched at the neural plate border consistent with a role for ALK in the delaminating neural crest cells (Figure 6A, 6D). Furthermore, ALK continues to be expressed at 9.5dpc at the neural plate border, and in a migratory neural crest destined for branchial arch I and II and at the frontonasal process (Fig 6D). Additional expression was seen in the heart, trunk and limbs. We also examined localization of the active form of ALK protein. Using an antibody recognizing ALK carrying a phospho-tyrosine residue (pY1507-ALK), we again found enrichment of ALK in the right place at the right time to be acting upon GSK3 during neural crest migration (Fig 6B-F). This neural crest specific expression was recapitulated in explant cultures, where we noted a lack of ALK protein in neural plates (total ALK, Fig 6G-G’) followed by an onset of expression in migratory neural crest cells, which was somewhat nuclear, with diffuse staining throughout the cell (compare Figure 6H-I’).

**Figure 6.**
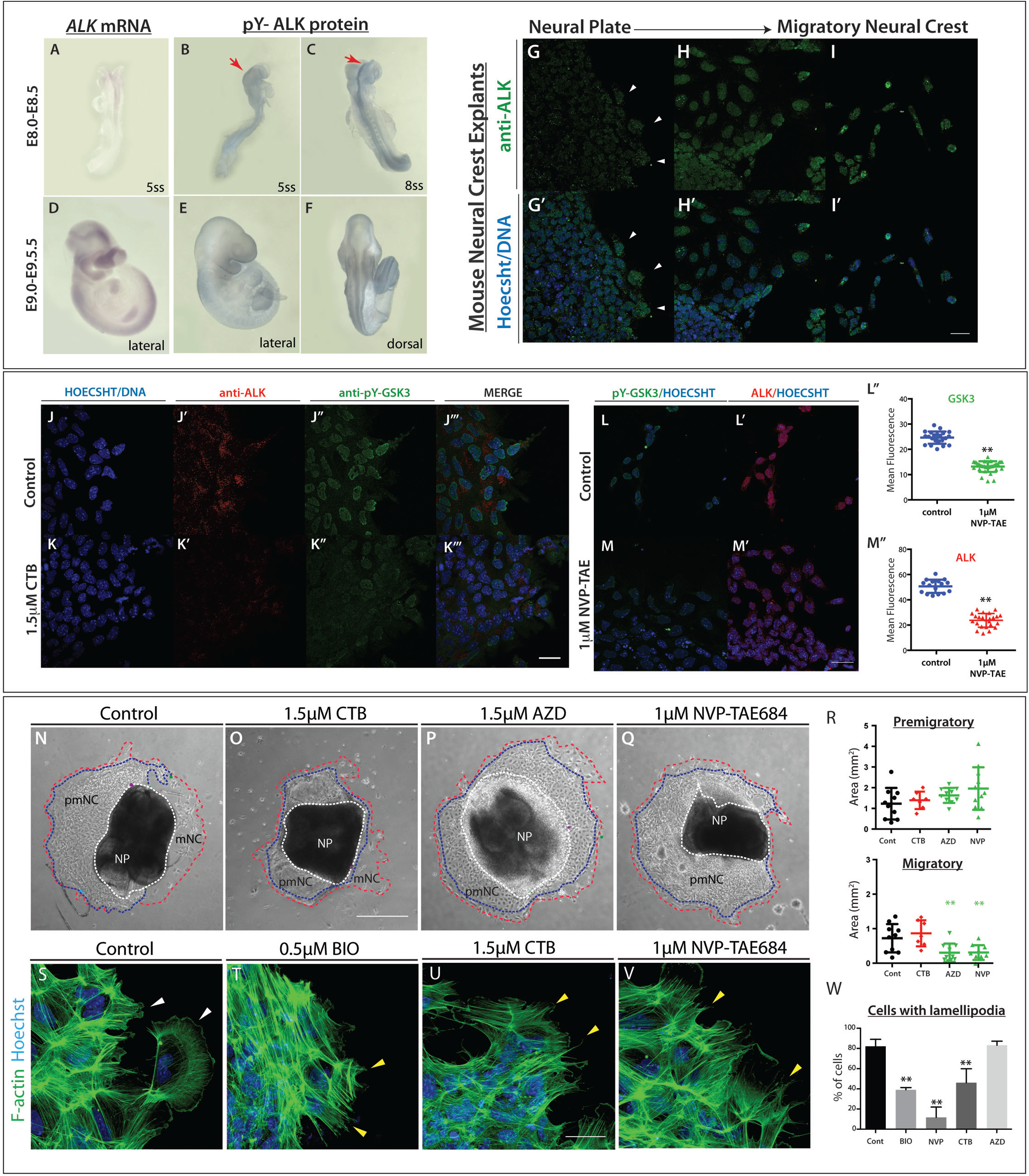
Inhibition of anaplastic lymphoma kinase leads to decreased levels of pY-GSK3 in mouse neural crest. (**A-F**) Anaplastic lymphoma kinase (ALK) is expressed in the neural crest. (A, D) mRNA *in situ* hybridization for *Alk* in e8.5 and e9.5 mouse embryos. (B-C, E-F) Antibody staining for activated ALK protein shows expression at the neural plate border (red arrows) and in the branchial arches. (G-I’) Staining for total ALK (green). (G) Very little total ALK is present in the neural plate with some present at the edge of the premigratory neural crest (white arrowheads). (H) Migratory neural crest cells express higher levels of ALK. (I) In fully migratory NC cells ALK appears to be nuclear. (G’-I’) Anti-ALK merged with Hoechst. (J-J’’’) Co-immunostaining of pY-GSK3 (green) and ALK (red) show that ALK is expressed in all cells that express pY-GSK3. (K-K’’’) Treatment with the ALK inhibitor crizotinib (CTB) for 24 hours reduces the levels of pY-GSK3. (L’’ and M’’) Quantitation of loss of pY-GSK3 and ALK fluorescence using the alternative inhibitor NVP-TAE also results in a loss of mean fluorescence intensity. (N-Q) Bright field images of neural crest explants treated with vehicle control or three different ALK inhibitors: 1.5*μ*M crizotinib, 1.5*μ*M AZD-3463, or 1*μ*M NVP-TAE-684. All three treatments showed a loss of migratory neural crest (red dotted lines.) (R) Area quantification of the premigratory neural crest (pmNC, area depicted by blue dotted lines covered in N-Q) and the migratory neural crest population (mNC, red dotted lines). A significant reduction in mNC was seen in AZD and NVP treated explants. (S-V) Phalloidin staining shows F-actin structure in explants. Hoechst marks nuclei. Note the loss of lamellipodial structures upon ALK inhibition (U-V) is comparable to BIO treatment (T). (W) Percentage of cells with lamellipodial formations at the leading edge upon treatment with GSK3 inhibitor BIO or with ALK inhibitors.

### Inhibition of ALK activity leads to a loss of pY-GSK3 and phenocopies GSK3 inhibition

To test whether ALK activity is required for tyrosine phosphorylation of GSK3, we challenged the neural crest explants with specific inhibitors for ALK. These included crizotinib (CTB), which is currently used as a chemotherapeutic, and AZD3463 (AZD). Because both of these are dual function inhibitors (CTB blocking ALK and the c-met receptor^51,52^, and AZD blocking ALK and insulin-like growth factor receptors^53^), we also treated with the highly selective inhibitor NVP-TAE684 (NVP^54^) (Fig 6L-M). we found that blocking ALK led to a loss of pY-GSK3 expression in neural crest explants, suggesting that GSK3 was indeed a target of ALK kinase activity in this context (Fig 6J-M, L’’). Furthermore, in all three cases, blocking ALK function phenocopied loss of GSK3 activity leading to a substantial decrease in delamination of the neural crest and a loss of the migratory cell population (Figure 5N-Q). We noted that NVP treatment was the most effective at blocking neural crest migration while CTB had a much milder effect (Fig 6R). Finally, ALK inhibitors CTB and NVP phenocopied the disruption of the actin cytoskeleton seen when GSK3 was blocked (Fig 6S-W).

### Neuroblastoma lines with high levels of ALK also express high levels of pY-GSK3

We then wondered whether high levels of ALK activity could drive excessive activation of GSK3. To test this, we examined nine neuroblastoma lines and found a clear correlation between levels of total ALK, active (pY-1507) ALK and pY-GSK3 (Figure 7A). We then focused on the Kelly neuroblastoma line, which carries an activating mutation in ALK (F1174L)^55^ and, as a comparison, LS^56^ neuroblastoma cells which had very low levels of ALK. In western blots, we again found much higher levels of ALK in Kelly cells, with nearly undetectable levels in LS cells, more similar to that of mouse embryonic fibroblasts (MEFs) (Figure 7B).

**Figure 7.**
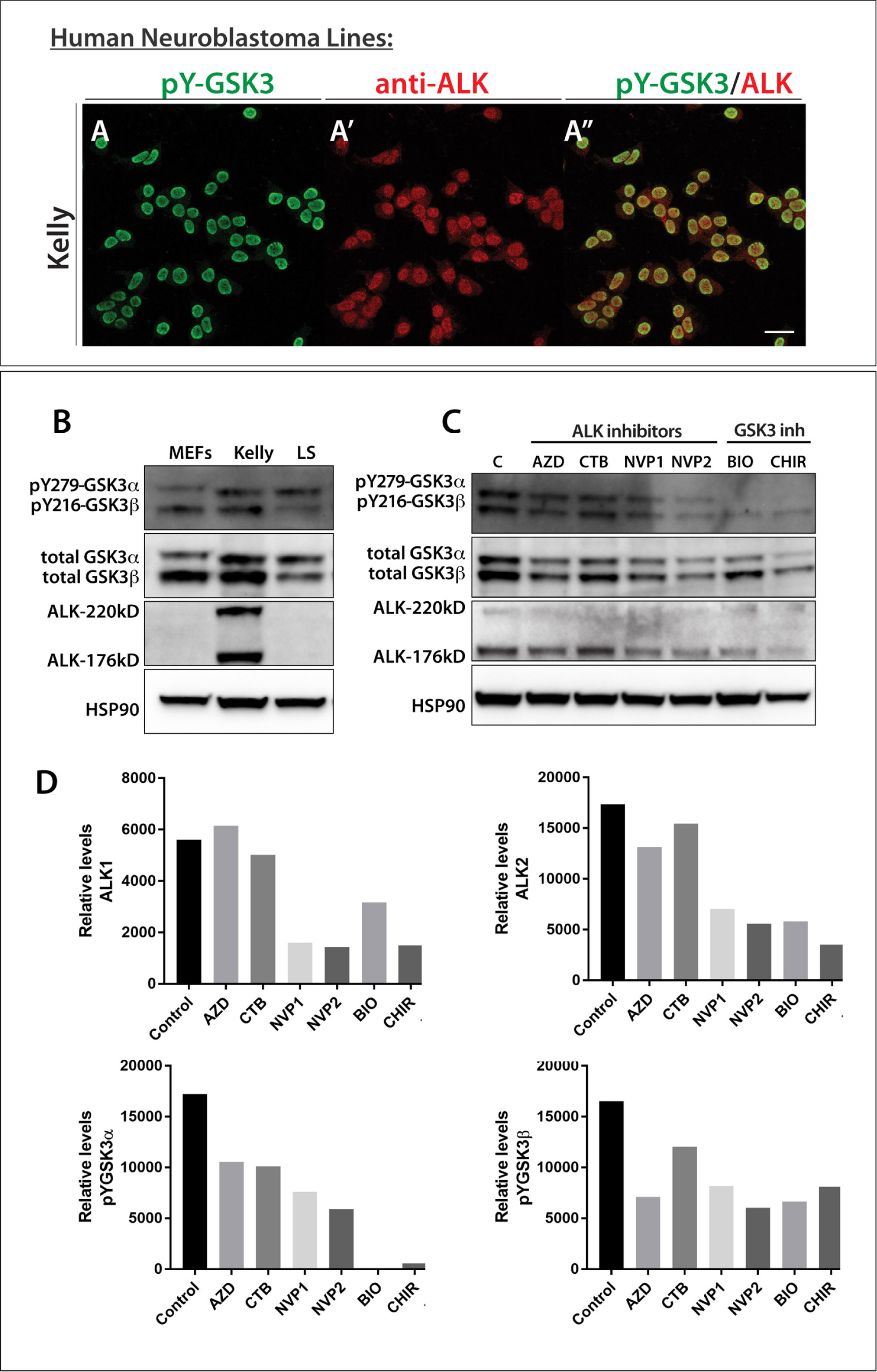
Neuroblastoma lines with high levels of active ALK also have high levels of pY-GSK3. (A) Western blotting of neuroblastoma lines reveals levels of pY-GSK3 correlates with levels of ALK-pY1507 (IMR5, Kelly, Be(2)C, IMR32, SH-SY-5Y, SK-N-SH). Cell lines with low or no ALK-pY1507 (SK-N-AS, LS, CHP-212) have correspondingly low levels of pY-GSK3, (B) Western blotting showing that only Kelly cells have high levels of ALK when compared to mouse embryonic fibroblasts (MEFs) and LS neuroblastoma cells. (C) Western blotting analysis reveals that, in Kelly cell line, ALK inhibition with NVP-TAE, results in gradual loss pYGSK-3α and pYGSK-3β are significantly reduced after 24h treatment with NVP-TAE, GSK3 inhibition, using BIO or CHIR, leads to a reduced expression of pY-GSK3, more predominantly is pY-GSK3α isoform. Some loss of ALK is also seen in NVP treatments. (D) Left, quantification of ALK (220kD) levels normalized to HSP90. Right, quantification of ALK (140 kD) levels normalized to HSP90. (E) Left, quantification of GSK3α levels normalized to HSP90; right, quantification of GSK3β levels normalized to HSP90. Treatments used were 1.5*μ*M crizotinib, 1.5pM AZD-3463, 1*μ*M NVP-TAE-684 (NVP1), 2*μ*M NVP-TAE-684 (NVP2), 0.5*μ*M BIO, and 1.0*μ*M CHIR99021.

### ALK activity is required for pY-GSK3 in neuroblastoma lines

All together, our data raised the possibility that ALK regulation of GSK3 is a neural crest specific activity that may have been co-opted during cancer progression. Indeed, inhibition of ALK in the neuroblastoma lines also decreased pY-GSK3 levels (Figure 7C-D, Supplemental Figure 6). The Kelly neuroblastoma line carries an ALK-F1174L mutation which renders it somewhat insensitive to the ALK inhibitor crizotinib (CTB)^55^. Therefore, as with the neural crest explants, we confirmed these findings using the two other inhibitors AZD and NVP (Figure 7D-E). We found that treatment with ALK inhibitors was sufficient to decrease phosphorylation of GSK3 (Fig 7E). Treatment with BIO or CHIR also affected pY-GSK3 levels, consistent with some auto-regulation by GSK3 itself (Fig 7D).

Finally, we set out to determine whether the excessive levels of pY-GSK3 could underlie the aggressive nature of neuroblastomas. If GSK3 activity is downstream of ALK in this context, we would predict that inhibition of GSK3 in Kelly cells would be sufficient to limit cell migration. Indeed, using scratch assays where we measured the movement of cells, we observed that Kelly cell migration was blocked by ALK inhibitors (NVP/AZD) similarly to GSK3 inhibition (BIO/CHIR) (Figure 8A, C,. As noted before, Kelly cells were resistant to CTB (Figure 8A, C,).

**Figure 8.**
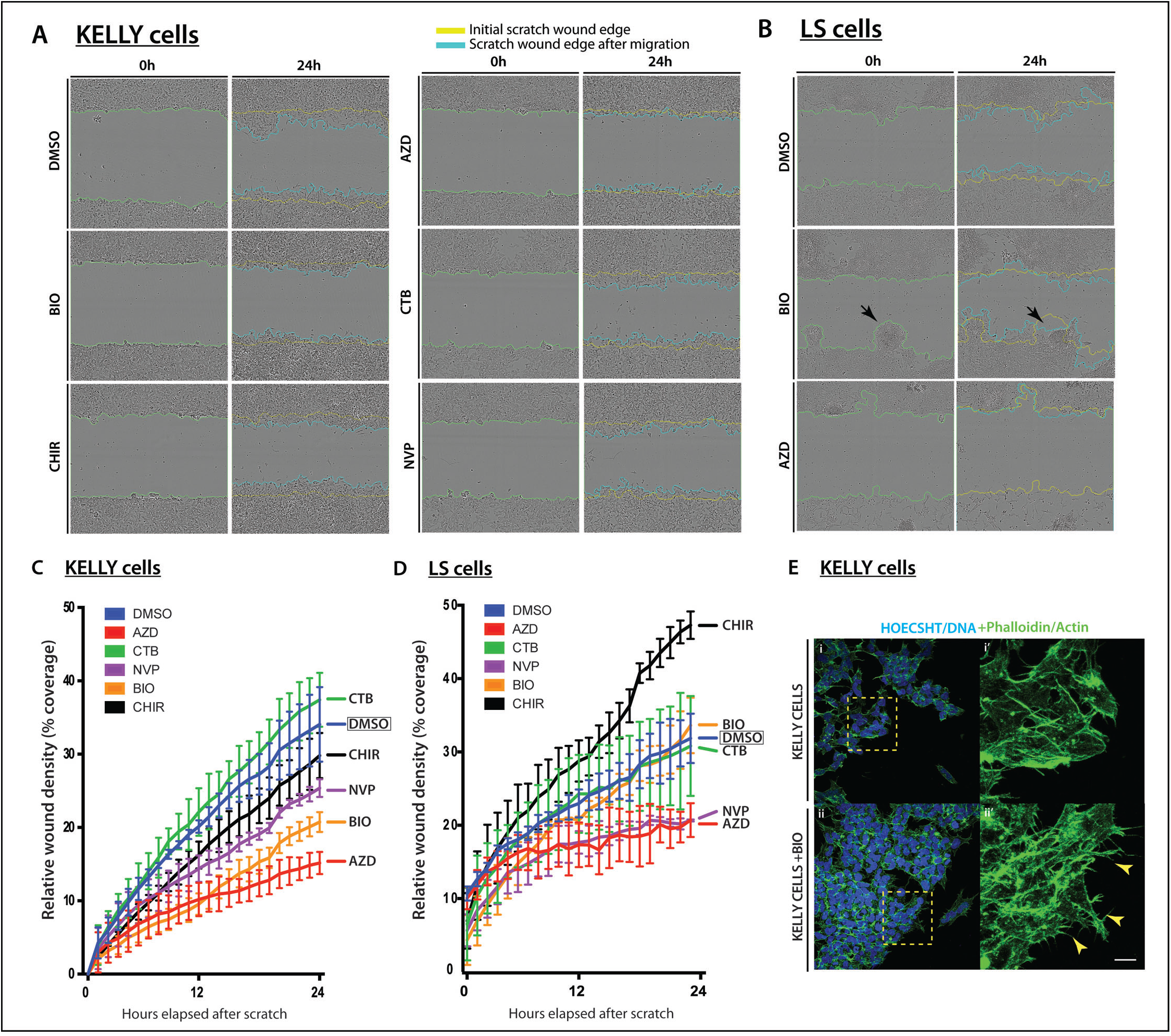
GSK3 and ALK inhibition affect migration in neuroblastoma cell lines. (A, C) Cell migration assay for Kelly neuroblastoma cell line. (A) Representative bright field still images at start (t=0h) and end (t=24h) time points of the migration assay in Kelly cells under various GSK3 (0.5μM BIO, 1.0μM CHIR99021) and ALK inhibition treatments (1.5μM CTB, 1.5μM AZD-3463 and 1.0μM NVP-TAE684). (C) Line graph representing the percentage of wound coverage over time. Notice that upon NVP-TAE684 (NVP) and BIO treatments cells do not close the wound as quickly as the control or unaffected CTB samples. Surprisingly AZD treatment showed the lowest percentage of wound coverage. (B, D) Cell migration assay for LS neuroblastoma cell line. (B) Representative bright field still images at start (t=0h) and end time (t=24h) points of the migration assay in LS cells under GSK3 inhibition (0.5μM BIO) and ALK inhibition treatment (1.5μM AZD-3463). Note that upon BIO treatment LS cells tend to aggregate and expand into the wound (black arrows). (D) Line graph representing the percentage of wound coverage in LS cells. There is a tendency to increase migration upon GSK3 inhibition (BIO and CHIR). The lower wound coverage upon ALK inhibition treatment correlates with reduced population of cells at the end of the assay suggesting a compromise in cell viability under these conditions. (E) Representative images of actin staining (phalloidin, green) showing Kelly cells treated with BIO compared to controls. Notice the irregular spiky protrusions of cells where GSK3 is inhibited (yellow arrowheads). Scale bar, 25 μm.

In contrast to the Kelly cells, LS cells, which do not carry the ALK-F1174L variant, behaved very differently. LS cells had substantially less pY-GSK3, which correlated with much lower levels of active ALK (pY1507, Figure 7A or pY1586, Supplemental Figure 6A). LS cells were insensitive to crizotinib treatment (Figure 8B, D and Supplemental Figure 6B-D). But, while the other ALK inhibitors led to a substantial decrease in the area covered, examination of the cultures showed substantial cell death (see Figure 8B, bottom right panel). More interesting, we found that GSK3 inhibition in LS cells elicited an unusual phenotype in the scratch assays, with cells moving together in aggregates, rather than as single cells (Figure 8B, arrowheads). It is possible that LS cells have a more “epithelial” morphology than Kelly cells and that GSK3 loss mimicked the stable cell-cell interactions similar to those in pre-migratory neural crest cells (Figure 3U-Y). Consistent with our hypotheses, morphologically, the Kelly cells responded similarly to motile neural crest cells, with BIO treated Kelly cells appearing compacted with dense actin staining (Fig 8E). Nevertheless, taken together, our data suggest that ALK activity is closely linked to GSK3 phosphorylation and activity in primary neural crest and neuroblastoma cells.

## Discussion

A key defining feature of the neural crest lineage is the ability to undergo EMT, acquire motility, migrate, and, ultimately, to differentiate into diverse cell types during embryonic development. Aberrant NC development results in neurocristopathies and other malignancies such as cancer. Thus, the same migratory characteristics could contribute to tumorigenesis and metastasis. The understanding of cellular behaviours in the normal context can aid our identification of important molecules involved in abnormal cell behaviours.

Here, we studied the effect of the serine/threonine kinase GSK3 during mammalian neural crest migration. Previously, we have found that GSK3β is required for the palate formation at specific time points during development^6^. This structure depends upon neural crest migration. However, due to functional redundancy between GSK3α and GSK3β, the *in vivo* activity has been difficult to study^57^. Furthermore, early loss of GSK3 leads to global Wnt activation, which can obscure later developmental roles^24,25,58–60^. However, our studies bypass these early complications and provide a refined understanding of the regulation and function of GSK3. This effect on neural crest migration appears to be β-catenin independent, as neural crest specific expression of stabilized β-catenin does not disrupt migration^61^. Instead, GSK3 appears to act directly on the actin cytoskeleton, changing the dynamics of lamellipodial formation. Our data demonstrate that this regulation may be via regulation of FAK localization, as well as key downstream factors including Rac1, cdc42 and lamellipodin. Interestingly, neural crest specific deletion of *FAK, Rac1, cdc42* and *lpd* all have a range of craniofacial anomalies^62,63^. However, it is worth noting that in our assays, we predominantly found that these proteins were relocalised, and thus it is difficult to directly compare our observations with the published null phenotypes. Nevertheless, these observations are worth further in-depth study.

GSK3 can also regulate the dynamics of the actin cytoskeleton, microtubules and cell to matrix adhesions^64^. To date, polarized inhibition via phosphorylation of S9 of GSK3β has been thought to be the main mechanism for establishment of cell polarity, especially in astrocytes^32^, and is also critical for glioma cell invasion^65^. All of these scenarios depend on negative regulation of GSK3. Importantly, contrary to the neuronal cell scenario, we find GSK3 inhibition via serine phosphorylation is not necessary for neural crest migration (Supplementary Figure 4). However, given the complexity of GSK3 regulation, it would be interesting to see whether combining phosphorylation mutations on the activating tyrosines and inhibitory serines has an additive effect on neural crest migration.

Most important, we find that neural crest cell migration depends on GSK3 activity, and that this correlates with high levels of tyrosine phosphorylation via ALK. While there are other kinases which may be regulating GSK3 phosphorylation, including GSK3 itself^16^, the association with ALK in the context of neuroblastoma is particularly compelling. However, future studies should include other tyrosine kinases which may be phosphorylating GSK3. For instance, it has been reported that PYK2^66^, a putative mammalian homologue of ZAK1, a kinase found in *Dictyostelium* shown to phosphorylate GSK3β at Y216. However, there is still no clear evidence on how this finding could relate to mammals. In pathological conditions, pY216-GSK3β has been found in prostate cancer, and Src was found to promote this phosphorylation, and with it, cancer progression and invasion^67^. These other kinases are worth considering in the future, and may be necessary to regulate sub-populations of GSK3.

Particularly intriguing was the prediction that GSK3α is a putative ALK substrate in cancer cells^18^. ALK is negatively correlated to neuroblastoma prognosis, with hyperactivating mutations of this kinase found in some of these aggressive tumors. Therefore, the discovery that pY-GSK3 was specifically expressed in delaminating and migratory neural crest cells, and that this correlated with ALK-positive neuroblastoma cells was extremely exciting. The additional novel discovery that ALK inhibition perturbs neural crest migration as well as expression of pY-GSK3 provides strong evidence for a new signaling cascade regulating neural crest cellular behaviour. Finally, because we focused on cell motility and lamellipodia formation, we cannot exclude the possibility that the ALK-GSK3 pathway has additional or longer-term effects on neural crest differentiation. Neuroblastoma lines carrying ALK-F1174L have also has been suggested to regulate serine (S9) phosphorylation of GSK3β via activation of the PI3K/AKT pathway. This leads to the stabilization of MYCN resulting in the formation of aggressive, highly penetrant tumours^45^. Because this serine phosphorylation was not required in the endogenous neural crest, we did not address the status of inhibited (phospho-serine) GSK3 in neuroblastoma cells. Nevertheless, this raises the possibility that phosphorylation events are necessary to set up distinct pools of GSK3 within the cell, which then regulate cell motility, transcriptional activity or protein localisation.

All together, our study demonstrates that the timing and control of active GSK3 within the embryo plays important roles during multiple steps in neural crest development. These lessons from the embryo improve our understanding of the pathological misregulation of key kinases, in neuroblastoma and congenital anomalies, and may have broader implications for cell motility in diverse systems.

## Materials and Methods

### Animal Procedures

All animal work was performed at King’s College London in accordance with UK Home Office Project License 70/7441 (KJL).

Mouse strains: CD-1 mice were obtained from Charles River Laboratories. *GSK3α^lacZ^, GSK3β^lacZ^, GSK3α^fl^* and *GSK3β^fl^* mouse lines have all been described previously^57,68^. The following Cre drivers were used: *pCAGG::Cre-ER^t2^* ^43^, *Wnt1-cre: Tg(Wnt1-cre)11Rth*^28^. The following reporter lines were used: *R26R^mT-mG^: GT*(*Rosa*)*R26Sor^Tm4(ACTB-tdTomato-EGFP)Luo^* ^69^. Genotyping was performed as described in original publications.

Mouse handling: The gestational ages were determined based on the observation of vaginal plugs which was considered embryonic day 0.5 (E0.5). Embryos were further staged by counting the number of somites after dissection. For each experiments, a minimum of three mutants with litter-matched controls were studied unless otherwise noted.

*Xenopus laevis:* Embryos were cultured using standard methods^70^. Staging was according to Nieuwkoop and Faber^71^.

### Cell culture

Mouse embryonic fibroblasts (MEFs) were prepared according to standard procedures and cultured in Dulbecco’s modified Eagle’s medium (DMEM, Sigma) supplemented with 10% (v/v) fetal bovine serum (FBS), 2mM L-glutamine, Pen/Strep, 15mM HEPES, b-mercaptoethanol (all from Invitrogen). Neuroblastoma (NB) cell lines LS and Kelly were cultured in RPMI media supplemented with 10% (v/v) FBS and Pen/Strep. For tamoxifen dependent cre deletions in culture, 4-OH-tamoxifen (Sigma H7904-5mg) was applied at 2μg/ml for 24h. After the incubation period, the media was replaced with standard media and cultured at 37°C and 5.0% CO_2_. Mouse primary cranial neural crest cultures were performed according to^72^.

For cranial mouse neural crest explants, dissections were performed on embryos at 8.5 days post coitum (dpc) and only embryos at the 5-8 somite stages were used. Briefly, the embryo was positioned to visualize and excise the neural fold. Adjacent tissues, such as mesoderm, were carefully cleaned from the neural plate. The neural plate was then cultured on matrigel-coated plates or slides in basal neural crest media at 37°, 5.0% CO_2_ for 24h to allow migration of neural crest cells out of the NP. When drug treatment was applied, it was added at plating (T=0).

Neuroblastoma cell lines used are noted followed by the source. All cell lines were mycoplasma tested. <u>LS^56,73^, DMSZ (ACC 675) lot #1; CHP-212^74^, ATCC CRL-2273 lot #58063161; Kelly^75,76^, DMSZ (ACC 355) lot #7; SK-N-SH^77^, HPA lot #09/D/005; SH-SY-5Y^78,79^, DMSZ (ACC 209) lot #12; IMR32^80,81^, HPA lot #04/K/012; BE(2)C^82^, ATCC CRL-2268 lot #10H023; SK-N-AS^83^, ATCC CRL-2137 lot #58078525; and IMR5^84^, gift from Martin Eilers, Wurzburg</u>.

### Antibodies

Primary antibodies used for immunofluorescence (IF) or Western blotting (WB):

mouse phospho GSK3 Tyr279/216 (Millipore 05-413, 1:300 IF; 1:1000 WB) rabbit GSK3b (Cell Signaling #9336) rabbit p75NTR (Millipore 07-476, 1:500 IF) rabbit total ALK (D5F3 Cell Signaling Tech #3633, 1:300 IF) rabbit phospho-Y1507 ALK (Cell Signaling Tech #14678, 1:200 IHC) rabbit pFAK (Abcam 39967, 1:300 IF; 1:1000 WB) rabbit RAC1 (Santa Cruz SC-217, 1:500 IF) mouse CDC42 (Santa Cruz SC-8401, 1:300 IF) rabbit lamellipodin (provided by Krause lab, 1:200 IF) rabbit GAPDH (Cell Signaling #2118) mouse HSP90a/b (Santa Cruz SC-13119, 1:1000 WB) anti-muscle antibody (DHSB, 12/101, supernatant 1:5).

Secondary antibodies used:

mouse IgG-Alexa 488 (1:1000 IF) mouse IgG Alexa 568 (1:1000 IF) rabbit IgG-Alexa 488 (1:1000 IF) rabbit IgG-Alexa 568 (1:1000 IF) Mouse IgG Peroxidase (1:10,000 WB) Rabbit IgG-Peroxidase (1:10,000 WB)

### Immunoblotting

Cells were washed twice with phosphate-buffered saline (PBS) and lysed with RIPA lysis buffer with added phosphatase inhibitor (PhosStop, Roche) and protease inhibitor (cOmplete™, Roche). Protein concentrations were determined by Bradford protein assay (BioRad). Proteins were resolved by SDS-PAGE on 4-12% precast gels (NuPAGE^®^Bis-Tris), then transferred to PVDF membrane (IPVH00010, Millipore) using Trans-Blot^®^ SD semi-dry transfer cell (BioRad). Immunoblots were developed using chemiluminescent HRP substrate (ECL) (Immobilon, Millipore) and ChemiDoc™ Touch Imaging System (BioRad) for detection. Densitometry was performed using Fiji-ImageJ analysis software and all the values were normalized to HSP-90 control^85^.

### Embryo fixation and histology

Mouse embryos at e10.5 were collected in cold PBS and fixed overnight in 4% paraformaldehyde (PFA) at 4°C. Mouse embryos at e8.5, e8.75 or e9.5 were collected in cold PBS and fixed in 4% paraformaldehyde for 2h at 4°C.

Samples were subsequently washed in PBS three times at room temperature for 10 min each time then treated as corresponding subsequent method. For sections, samples were cryoprotected by incubating at 4°C overnight in graded sucrose solutions in PBS. Tissues were then embedded in OCT and frozen at -20°C. Heads and bodies were sectioned at 10μm and frozen at -20°C after drying at RT for 1h. For whole mount mRNA *in situ* hybridization, β-galactosidase activity and immunohistochemistry, samples were treated according to standard procedures.

*Xenopus* embryos were collected at the indicated stages and fixed for 1 hour in MEMFA at room temperature, before washing into ethanol for storage.

*ALK* cDNA clone was obtained from Source Biosystems (ID D130039F03). *Sox10* cDNA was a gift of the Pachnis lab^86^. *Life-ActGFP* constructs were a gift of the Mayor lab^41^.

### *Xenopus* cartilage staining

Stage 45+ embryos were fixed in MEMFA for 1 hour at room temperature before washing into ethanol. For cartilage staining embryos were washed into a 0.15% alcian blue solution (70% EtOH/30% acetic acid) at room temperature for 3 days. When suitably stained, embryos were rinsed 3x 15 mins in 95% EtOH. Rehydration was done stepwise into 2% KOH then washed from 2%KOH stepwise into 80% glycerol/20% 2%KOH, 1 hour per wash before washing overnight into the final solution. Dissection of cartilages was then performed to increase visibility of craniofacial cartilages.

### Dissection of *Xenopus laevis* neural crest and grafts

2-cell stage embryos were microinjected with membrane GFP or lifeact-GFP and then cultured at 15°C until they reached an appropriate stage. The procedure followed to obtain clean neural crest was as Milet and Monsoro-Burq^87^. Neural crest explants were plated in fibronectin-covered slides to study in vitro migration. They were incubated under control and 0.5mM BIO at time=0. Explants were examined 8h to 24h later, as indicated. For whole mount and grafts, the embryos were incubated in control and 15mM BIO and collected when they reached the desired stage.

### Drug treatments

All drugs were prepared at the concentration indicated, in the corresponding standard media for each cell type. GSK3 inhibitor 6-Bromoindirubin-3’-oxime, BIO (Calbiochem, 361550) was re-suspended in a stock solution of 140mM in DMSO and stored at -20°C until use for either mouse or *Xenopus laevis* experiments. The ALK inhibitor crizotinib (CTB) was re-suspended in a stock solution of 5.5mM in DMSO. The ALK inhibitor AZD3463 (Selleckchem, S7106) was re-suspended in a stock solution of 20mM in DMSO. All compounds were then further diluted in the appropriate media. For MEFs and NB cell lines, treatments were added when cells were at a confluence of at least 80%. Control treatments were performed at the corresponding DMSO concentration.

*Xenopus* embryos were incubated in 12-well plates, 20 embryos per well. For GSK3 inhibition 15 μM BIO (Calbiochem), was added to media or as otherwise indicated. Control embryos were incubated in 0.5% DMSO in media.

### Neuroblastoma cell scratch assay and single cell tracking

Cells were cultured to confluence in a 96-well ImageLock^TM^ plate (IncuCyte^TM^), in neuroblastoma media. At this point the cells were starved overnight in 2% FBS. For the scratch, a 96-pin mechanical device (WoundMaker™) was used to create homogeneous 700-800*μ*m wounds in the confluent monolayers after starvation. For detailed manual see IncuCyte^®^ Cell Migration Kit (Essen Bioscience). The following treatments, diluted in starvation media, were applied to the cells prior to the scratch making: 1.5*μ*M AZD-3463, 1.5*μ*M Crizotinib (CTB), 1.0 *μ*M NVP-TAE684, 0.5*μ*M BIO, 1.0*μ*M CHIR 99021 and DMSO as control. The plates were then incubated and scanned in the IncuCyte^®^ system at the rate of 1 scan/hour for up to 36 hours, but analysis was performed at 24-hour time point. Image processing and all the analysis were made using the IncuCyte^®^ ZOOM Software. Significance was based on two-tailed t-test.

### Microscopy and Image Analysis

Live imaging was obtained using Nikon A1R. Confocal z-stacks were obtained using a Leica TCS SP5 DM16000. Image sequences were reconstructed using Fiji-ImageJ analysis software.

---

**Supplementary Information** included below.

**Acknowledgements:** We thank J. Wallingford and M. Dionne for critical reading of the manuscript; M. Ishii, R. Maxson, P. Gordon-Weeks and Roberto Mayor for advice and help; and the NHH BSU and University of Portsmouth EXRC for animals. LC and co-workers are funded by Cancer Research UK and Children with Cancer UK. KJL and co-workers are funded by the Biotechnology and Biological Sciences Research Council, UK, the Medical Research Council, UK, CAPES Brazil and the King’s College London Dental Institute Newland Pedley Fund.

**Author Contributions:** SGGM, TB, DD and ALM designed experiments, collected and analyzed data. HAA collected and analysed data. MK, CP and LC provided crucial reagents. KJL designed the overall study, collected and analyzed data. SGGM and KJL wrote the paper with input from all other authors.

**Supplemental Figure 1.**
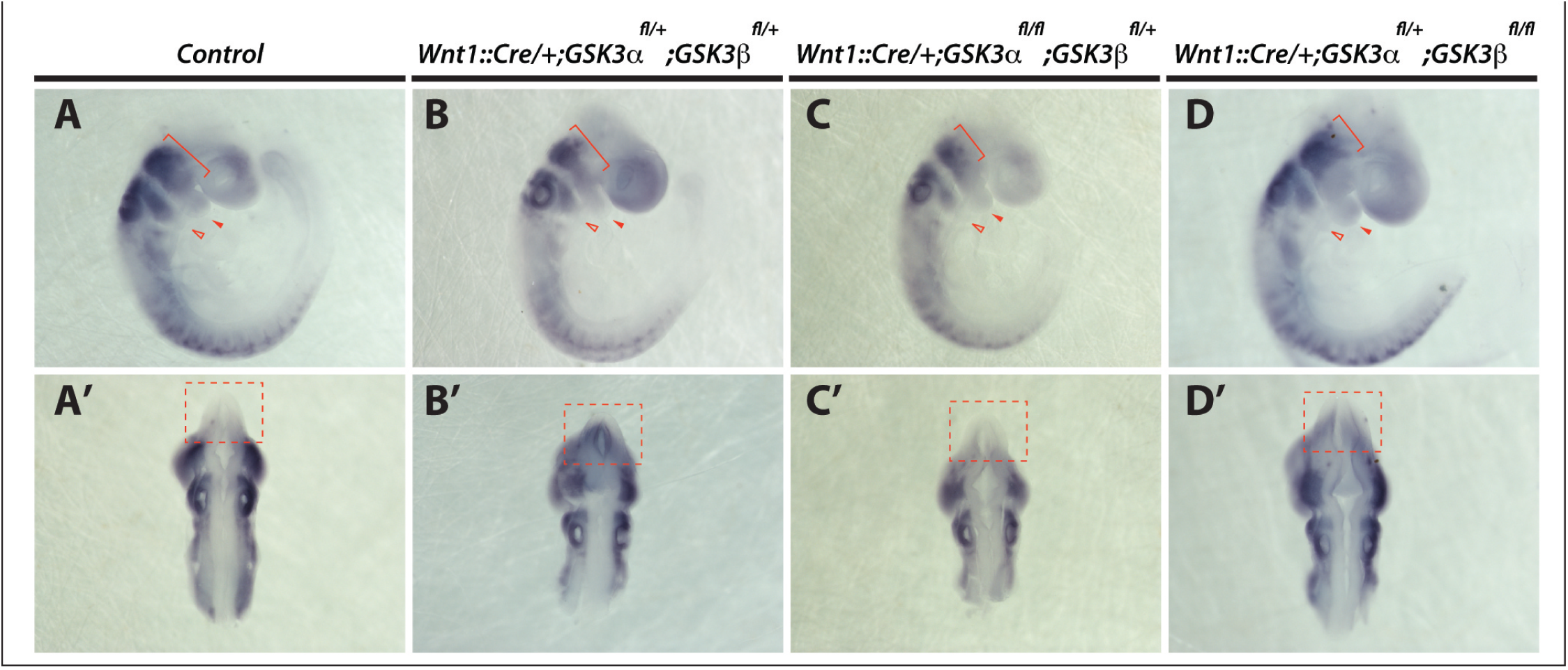
Neural crest migration shows subtle pattern differences depending on specific allele deletion of GSK3. (A-D’) Whole mount *in situ* hibridization for Sox10 labels migratory neural crest in E9.5 mouse embryos. (A-D) Lateral views. (A’-D’) Dorsal views. (A,A’) Sox 10 expression in control embryos is detected in the trigeminal, branchial arches, frontonasal process and the hindbrain neural crest streams at the level of r4-r6. Note absence of Sox10 positive cells in the brain (A’, red dotted square). (B,B’) Neural crest specific deletion of one allele of each GSK3 isoform results in accumulation of Sox10 expressing cells in the brain (B’, red dotted square) and slight reduced expression in branchial arch 1 (B, red arrowhead). (C,C’) Neural crest carrying only one allele of GSK3β showed subtle accumulation of Sox10 positive cells in the brain (C’, red dotted square), and reduced expression in branchial arch 2 and frontonasal process (red arrowheads). (D,D’) Neural crest specifically carrying only one allele of GSK3α, showed accumulation of Sox 10 positive cells in the brain (D’, red dotted square) and branchial arch 2 (red line arrowhead).

**Supplemental Figure 2.**
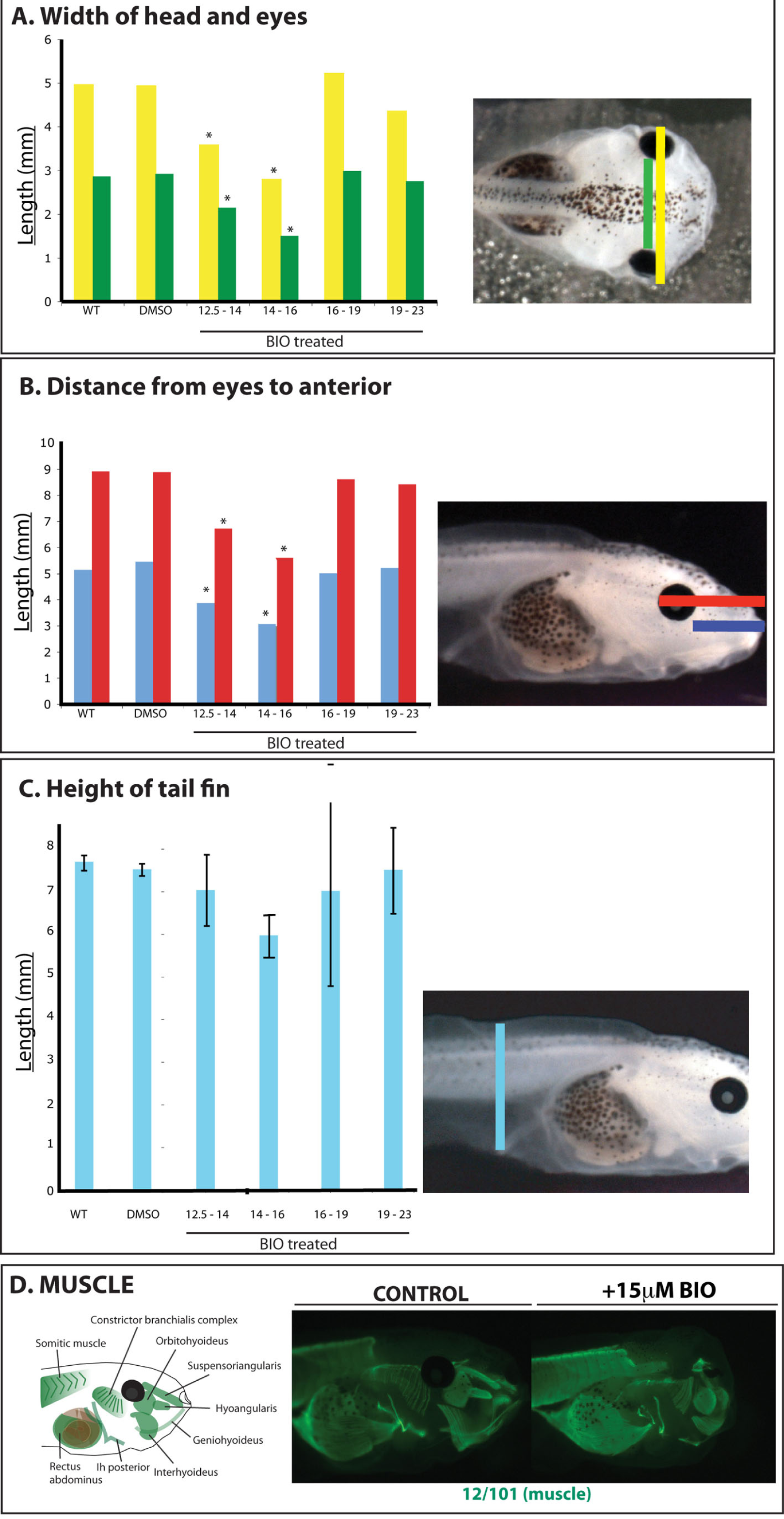
Timed GSK3 inhibition during frog (*Xenopus laevis*) neural crest migration results in embryo phenotypic defects. (A-C) Graphs showing the length measurement of different head structures obtained from st 45 embryos untreated and treated with GSK3 inhibitor, BIO. The timing of the treatment applied is specified in the X-axis of each graph. Note that treatments between st.12.5-14 and st.14-16, affect significantly head features such as the head width (A, yellow bars) and width between eyes (A, green bars), The same treatment timing has an effect on the lateral measurement of the distance measured from the posterior edge of the eye (B, red bar) and the anterior edge of the eye (B, blue bar) to the anterior edge of the embryo, (C) The height of the tail fin was found smaller in embryos treated from st.14 to st.16. (D) Schematic of craniofacial muscle in tadpoles. 12/101 antibody staining labeling muscle revealed that BIO treatment reduced size of craniofacial muscles (green) but does not perturb patterning.

**Supplemental Figure 3.**
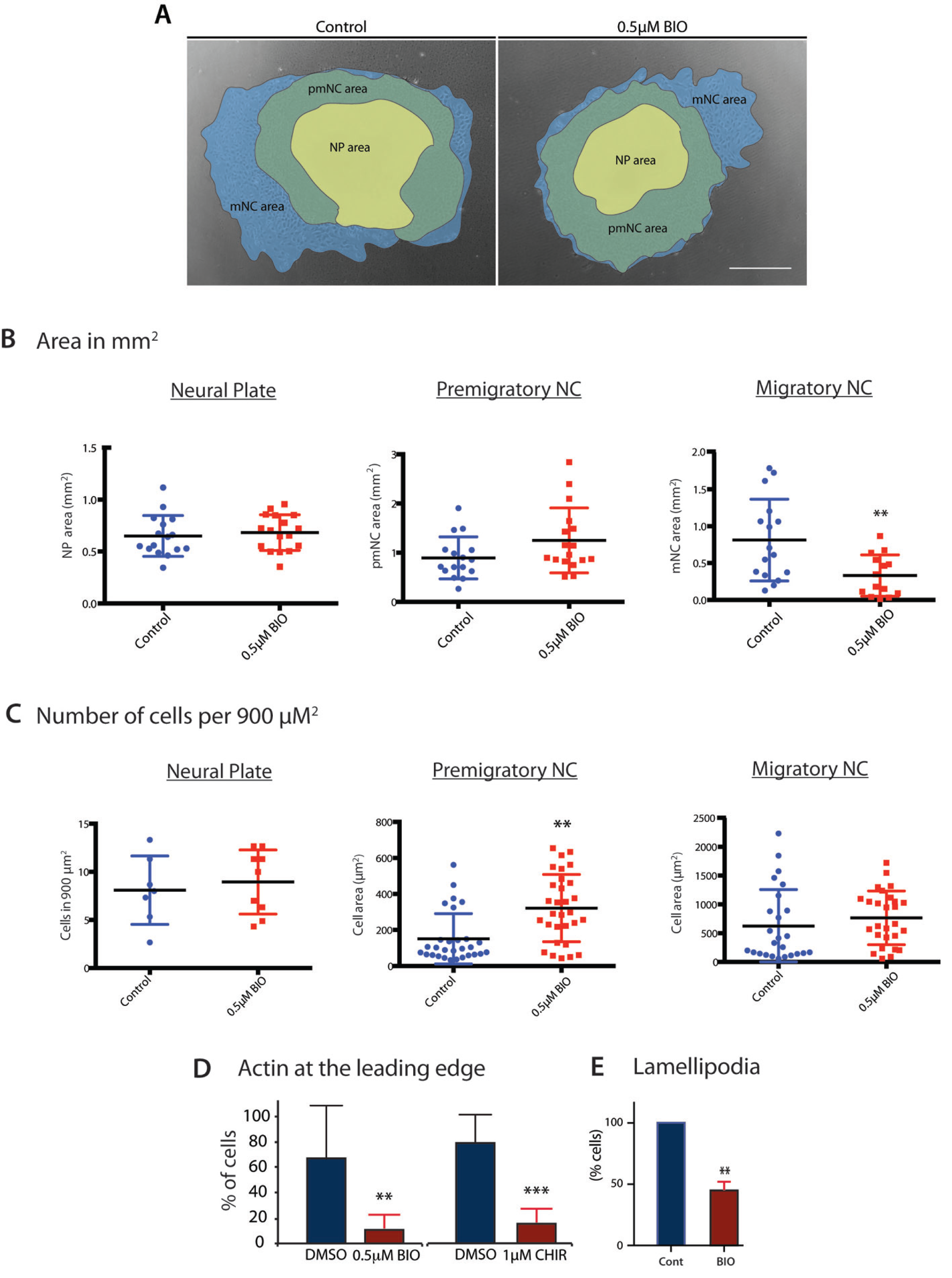
GSK3 inhibition reduces migration of neural crest cells, but does not affect the number or cell size of migratory cells. (A) Neural crest explants showing in different colour shades the cell populations described. In the middle the neural plate (NP), surrounded by a cell population that appears more epithelial, the pre-migratory neural crest (pmNC), and finally in the outer ring of the explant, the migratory neural crest (mNC) population where cells appear lose and show a mesenchymal phenotype. Neural crest cells treated with BIO, show a reduced expansion in the mNC. (B) Dot plots representing the absolute values obtained from the NP, pmNC and mNC areas. (C) Dot plots showing the number of cells contained in a specific area in control and BIO treated explants. Cell area in the pmNC appeared to be increased in BIO treated explants, however in the mNC population there was no difference between BIO treated and control samples.

**Supplemental Figure 5.**
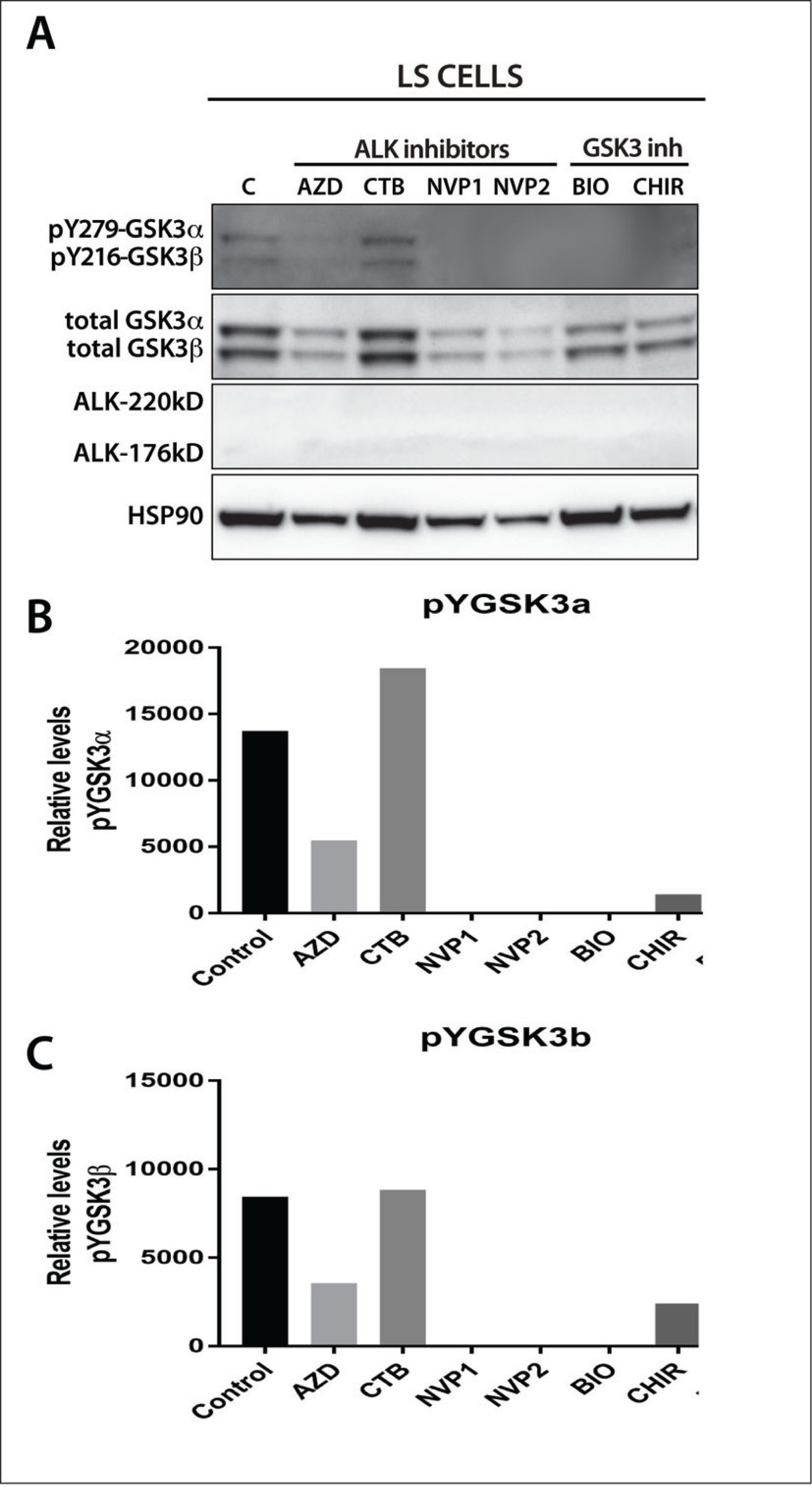
Serine 9/Serine 21 phosphorylation is not necessary for normal neural crest cell morphology and knockin mice are still sensitive to BIO treatment. (A) Phalloidin staining of neural crest explants from GSK3a^S21A/S21A^; GSK3b^S9A/S9A^ embryos shows actin at the lead edge of cells and in lamellipodia (white arrowheads). (B) Treatment with BIO results in more spiky filopodial protrustions and a loss of leading edge actin. Scale bar=20μM.

**Supplemental Figure 4.**
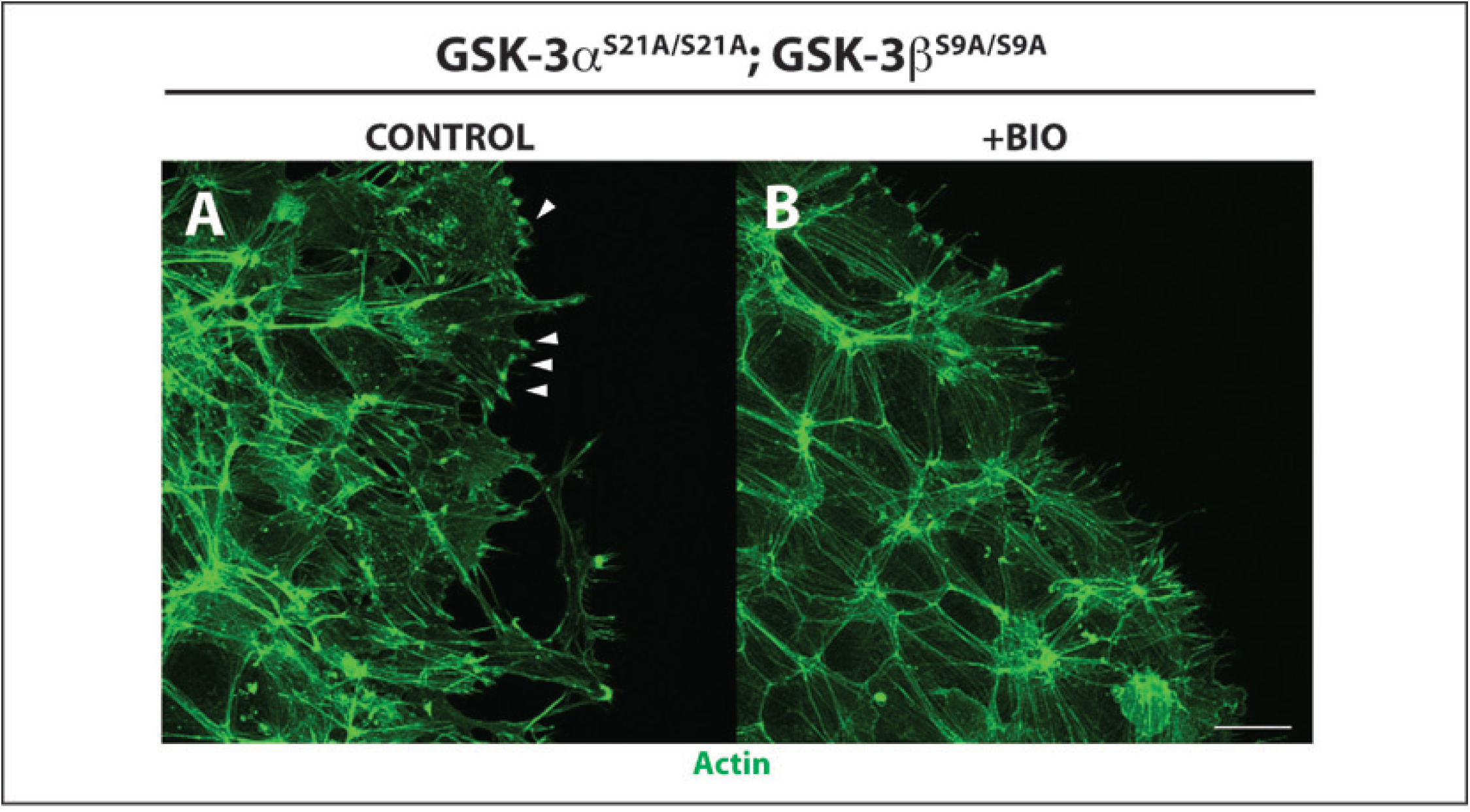
Effect of GSK3 inhibition on actin, lamellipodia and microtubules. (A) Bar chart showing the percentage of cells containing actin at the cell leading edge. GSK3 inhibition by two different compounds resulted in a significant reduction of this percentage. (B) Bar chart showing the percentage of cells that formed fan-shaped lamellipodia in control and BIO treated explants. (C) In controls (left) stabilised microtubules marked by acetylated α-tubulin are distributed throughout the cell. In BIO treated samples (right), acetylated α-tubulin staining is localized perinuclearly, with a bias towards the leading edge of the cell. Relative fluorescence is somewhat decreased through cell in treated explants. Schematic depicts relocalisation of acetylated α-tubulin staining. (D) In controls (left) unstable microtubules, marked by YL1/2, staining are distributed throughout the cell while in BIO treated samples YL1/2 staining is perinuclear and biased toward the posterior of the cell. Relative fluorescence is significantly decreased in BIO-treated samples (*p≤*0.05*). Schematic depicting relocalisation of YL1/2 staining.

**Supplemental Figure 6. pY-GSK3 profile in LS neuroblastoma cell line treated with ALK and GSK3 inhibitors.** (**A**) Mesoscale discovery (MSD) assay showing relative levels of active ALK (pY1586) and total ALK in the neuroblastoma lines shown in Figure 7A. Note low levels of ALK in LS cells. (B) LS cells were treated with ALK inhibitors (1.5μM CTB, 1.5μM AZD-3463 and 1.0μM NVP-TAE684) and with GSK3 inhibitors (0.5μM BIO, 1.0μM CHIR99021) for 24h and analysed by western blot for pY-GSK3, total GSK3 and ALK. Analysis confirmed absence of ALK in this cell line and a lower amount of pY-GSK3 than Kelly cell line (see Figure 7D). Treatment with ALK inhibitors showed that CTB did not affect the pY-GSK3 content compared to untreated cells, however AZD and NVP seem to have reduced total GSK3 and pY-GSK3 significantly, possibly due to loss of cell viability. GSK3 inhibitors maintained total-GSK3 unaffected, however pY-GSK3 expression was not detected. (C, D) Relative levels of pY-GSK3α (C) and pY-GSK3β (D) to loading control.

**Movie S1: Movie of control neural crest explants expressing *LifeAct-GFP*.**

*Xenopus* embryos were injected with mRNA encoding *LifeAct-GFP* at the two-cell stage. Neural crest explants were taken at stage 17 and cultured for 8 hours before imaging. This movie corresponds to Supplemental Figure 2E-H.

**Movie S2: Movie of BIO treated neural crest explants expressing *LifeAct-GFP*.**

*Xenopus* embryos were injected with mRNA encoding *LifeAct-GFP* at the two-cell stage. Neural crest explants were taken at stage 17 and cultured for 8 hours in 0.5*μ*M BIO before imaging. This movie corresponds to Supplemental Figure 2I-L.

**Movie S3: Movie of control mouse neural crest**

Movies of migrating neural crest from controls (*GSK3α^fl/fl^; GSK3β^fl/fl^; Rosa^mTmG/+^*) showing normal filopodial and lamellipodial dynamics, as well as migratory behaviour.

**Movie S4: Movie of GSK3 mouse mutant neural crest**

Movies of mutant neural crest explants (*pCAGG::CreER^tm^; GSK3α^f//fl^; GSK3β^fl/fl^; Rosa^mTmG/+^*) showing loss of motility and lamellipodial dynamics, but still showing filopodia formation.

**Movie S5: Merge of control and mutant neural crest from Movie S4 and Movie S5**

Experiments shown in Supplemental Movie 3 and 4 were performed in the same dish. Two sets of neural crest were plated together: *Cre* negative controls (*GSK3α^fl/fl^; GSK3β^fl/fl^; Rosa^mTmG/+^*) are labeled in red (membrane Tomato, mT) while *Cre* positive mutants (*pCAGG::CreER^tm^; GSK3α^fl/fl^; GSK3β^fl/fl^; Rosa^mTmG/+^*) are labeled in green (membrane GFP, mG).

**Movie S6: Brightfield movies of control mouse neural crest cells corresponding to Figure 5I.**

Movies of migrating neural crest from controls showing normal filopodial and lamellipodial dynamics, as well as migratory behaviour.

**Movie S7: Brightfield movies of mouse neural crest cells treated with BIO corresponding to Figure 5H.**

Movies of migrating neural crest from mouse explants treated with BIO showing loss of lamellipodial dynamics, as well as decreased cell movements.

